# On the Stability of Silicone-Encapsulated CMOS ICs for Active Implantable Devices: 4.3 Years of Accelerated Life Testing

**DOI:** 10.1101/2025.05.21.655306

**Authors:** Ahmad Shah Idil, Callum Lamont, Kambiz Nanbakhsh, Federico Mazza, Vasiliki Giagka, Tim Constandinou, Anne Vanhoestenberghe, Nick Donaldson

## Abstract

The reliability of polymer-encapsulated CMOS integrated circuits (ICs) is critical for the development of miniaturised active implantable medical devices (AIMDs). Traditional hermetic packaging methods become impractical as implant sizes decrease, necessitating alternative methods of protection from body fluids. This study evaluates the long-term stability of silicone-encapsulated CMOS ICs through accelerated life testing using electrical impedance spectroscopy (EIS) and visual inspection. CMOS interdigitated combs (IDCs) were encapsulated in medical grade silicone rubber, subjected to immersion in phosphate-buffered saline (PBS), and tested at elevated temperatures (47°C, 67°C, and 87°C) under both 5V DC and biphasic voltage biases for up to 4.3 years.

Remarkably, no insulation failures were observed in the IDCs, with no significant water ingress detected through impedance changes. Failures at the ICs were limited to wire bond open-circuits, though there was some pad discolouration/corrosion. Other failures were elsewhere, not at the ICs. This highlights the stability of modern silicon oxide/silicon nitride bilayer passivation when encapsulated in adhesive silicone rubber. Visual analysis revealed occasional solder and aluminium pad corrosion, particularly at higher temperatures, but these changes did not correlate with EIS failures. The findings suggest that silicone encapsulation, combined with passivation and shielding strategies, enables long-term IC reliability in biofluid environments.

To our knowledge, this paper presents the longest reported accelerated ageing study of test structures for implantable devices, laying the groundwork for the integration of silicone-encapsulated ICs into next-generation chip-scale bioelectronic implants.

## 1. Introduction

In almost all active implantable medical devices (AIMD), electronics are housed inside a gas-filled, hermetically-sealed titanium enclosure, following a method initially proposed by Cowdrey at Telectronics in 1971 [53]. Early attempts at polymer encapsulation (usually epoxy) to cover all component surfaces exhibited relatively poor reliability and were generally discarded [31]. Hermetic packaging maintains electronics, particularly regions of high field strengths, in a dry environment, thus achieving long implant lifetimes [60]. However, this approach becomes increasingly impractical as device sizes decrease. For small implants with electronics placed close to electrodes, as envisioned for bioelectronic medicines, metal enclosures are excessively bulky. Another approach is micropackaging, similar to methods used in microelectromechanical systems (MEMS), where a lid is bonded to the integrated circuit chip [51]. However, wafer-on-wafer bonding processes for these micropackages can be prohibitively expensive, particularly for university research, and as internal volumes shrink, the methods for validating hermeticity become inadequate [59].

Metal enclosures present other practical challenges, such as wireless data and power transmission, high-density interconnections for electrode arrays, sensor interfaces, and maintaining optical transparency for optogenetic implants [27,67]. The research presented in this paper emerged from the CANDO project, that aimed to chronically implant micro-LEDs in cortical tissue (Wellcome Trust WT102037, EPSRC NS/A000026/1) to control neural activity optogenetically. The project necessitated exploring alternative encapsulation methods to ensure device longevity in the body [27, 67].

An alternative is silicone encapsulation, which may suffice for protection. The Brindley sacral anterior root stimulator implant (SARS), commercialised by Finetech Medical, has demonstrated a well-established track record of long-term reliability with components encapsulated in silicone rubber (PDMS) [48] (see Figure 1). A review of the first 500 implants by Brindley reported mean times to failure of 19 years [10]. The mechanism by which silicone encapsulants protect implanted electronics is often misunderstood. Silicone does not function as a water vapour barrier akin to hermetic packaging; indeed, it is highly permeable to water [9]. The critical properties of silicone encapsulation include strong adhesion to surfaces, resistance to chemical attack, purity, and low mechanical modulus. Properly applied silicone encapsulation without voids prevents liquid condensation despite rapid saturation with water vapour, thus avoiding ionic current flow and subsequent corrosion [17, 19, 54, 60].

**Figure 1.**
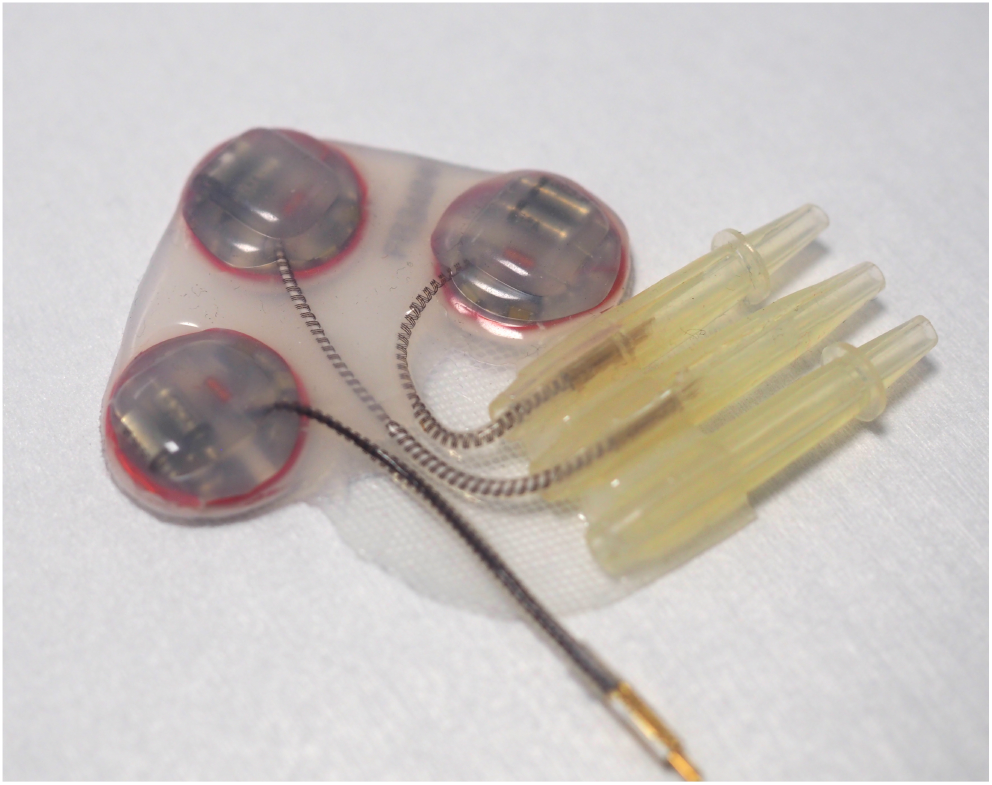
The silicone-encapsulated Finetech-Brindley SARS implant, with a mean time to failure of 19 years. It consists of only discrete components and no CMOS IC. Reproduced from [40], licensed under CC BY 4.0.

Given the predominance of complementary metal–oxide–semiconductor (CMOS) technology for integrated circuits that power medical devices, this study evaluates whether silicone-encapsulated CMOS ICs could reliably support chronic implants lasting beyond 28 days. To test this hypothesis, we developed and validated a dedicated system, the ALTA (“Accelerated Life Testing Apparatus”) which was used in this study to perform long-term accelerated ageing tests using electrical impedance spectroscopy (EIS). The design and validation of ALTA are detailed in a previous publication [16].

Initial studies focused on interdigitated comb structures with coarser geometries and various combinations of passivation materials and silicone encapsulants [41]. The promising results warranted further investigation using actual CMOS devices, which this paper presents.

The remarkable stability demonstrated herein, specifically the absence of insulation failures after 4.3 years of accelerated ageing under elevated temperature and continual voltage bias, motivated further companion investigations into passivation layer performance and potential encapsulation breach mechanisms [45]. As reported in that work, the comparable condition of the passivation layer in silicone-encapsulated ICs aged in vivo and in vitro in PBS lends confidence to the use of long-term in vitro studies as a valid proxy for implant conditions, thus reinforcing the promise of silicone encapsulation for next-generation bioelectronic implants.

### 1.1. Context and Paper Aim

Before undertaking the tests we report in this paper, our expectation was that integrated circuits in saline, protected only by silicone encapsulation, would fail by insulation breakdown due to the relatively high field strengths (MV/m) present, as the field strength had been established as an important parameter in our review of failure mechanisms [60].

In the CANDO project, we hypothesised that the use of LED-driving waveforms with symmetric positive and negative phases would reduce the rate of electrochemical degradation when ionic leakage currents started to flow. However, if the implant was to be active all the time, the power supply voltage would be present continuously, creating a “worst-case condition” with continuous bias (“DC”). Previous studies on parylene-coated platinum tracks have shown that continuous DC bias leads to more severe degradation, primarily through electrolysis-induced cracking, than symmetric biphasic signals, which instead result in gradual platinum dissolution and migration [44]. Hence, we needed to evaluate how long the devices would remain functional with either DC or biphasic bias (“ageing voltages”).

Our aim with these tests was to collect data to evaluate whether the device would function for a reasonably long time implanted in a human. We tested samples at three elevated temperatures because if the failure mechanism was the same at all three, this would enable us to extrapolate a sample lifetime at body temperature (see Section 5.5).

Because we did not expect long lifetimes in our accelerated ageing tests, as we had no knowledge of how CMOS IC passivation layers would survive in saline [45], we included in the design features that we believed would mitigate the destructive process: a protective metal “shield layer” above the IDC to prevent hydration of the dielectric layer directly above the IDC, and a “wall-of-vias”, in addition to the standard die seal ring, to protect the edges of the chip from delamination and lateral water penetration. The sample design and experimental setup are presented in Section 3.1.

The aim of this paper is to present the remarkable results of multi-year accelerated ageing tests, which, to the best of our knowledge, represent the longest duration reported for test structures intended for implantable devices. Contrary to our initial expectation, even after 4.3 years of immersion in saline, there were no insulation failures; the only IC failures observed were conduction failures (probably) from open-circuits at wire bonds.

## 2. State of the Knowledge

### 2.1. Purpose of Encapsulants

The problem of protecting electronics implanted in the body is an extreme case of the general problem facing users of integrated circuits operating at high humidity. Ionic currents flow in the moisture film that forms on every surface and whose thickness depends on the humidity above the surface. Encapsulants are used to prevent these surface currents. How they work was clearly stated by White in his 1969 paper Encapsulation of Integrated Circuits [63]:

> *“*…*[the] function of the coating in preventing leakage currents is to prevent moisture from adsorbing or condensing on the surface to form a continuous film which can act as a conducting path between metal conductors.”*

Many investigators have compared different polymers and have found low-modulus silicones most effective. A successful silicone encapsulation conforms to the surface’s topology to cover it without voids to prevent condensation on the surface. Corrosion and electromigration require a liquid film for the displacement of charges. Yan [65] plotted the evolution of surface conductivity over IDCs on alumina samples as monolayers of water adsorbed. If adsorption is uniform on a given surface and ignores chemisorption, a limit of three monolayers is often used [32].

The low Young’s modulus of silicone ensures low residual stress after cure, minimising potential delamination force. Excellent underwater stability, both of the polymer and of the adhesive interface, a property many adhesives do not have [49, 64], further ensures that even over time, no sites appear over the surface where liquid water could condense.

To address a common misunderstanding, we again stress that it is well-known that silicone rubber is not a good barrier to water vapour. In fact, it is highly permeable. Based on data from Lutz obtained for an unnamed methyl-vinyl based high temperature vulcanizing silicone rubber with aluminium hydroxide filler, a small void under a 5 mm thick layer of that silicone rubber, immersed in the body, would be saturated with water vapour within less than a minute [43]. Whilst there is a wide range of compositions of silicone rubbers, slightly affecting permeability to water vapour, the reason for their excellent performance in preventing degradation of electronics operating in high humidity environments is not that they act as a water vapour barrier but that they prevent condensation of water on the surface.

The metal tracks on integrated circuits (usually aluminium, copper, or an alloy) are protected by passivation layers that are deposited from chemical vapour (CVD). The passivation should be conformal, free of cracks and with low residual stress. Openings are formed over the pads to enable interconnection (wire bonds or otherwise). Because the pads and wire bonds must be protected, silicone is applied overall, so that the tracks are generally protected by both the passivation layer and the silicone encapsulant. Passivation is usually a layer of silicon oxide under a layer of silicon nitride. Silicone encapsulation insulates the connections to the bond pads (and perhaps other parts of the microelectronic assembly) and, as we have shown [45], plays a role in preventing dissolution of the passivation.

In this paper, we took the position of an IC designer wanting to maximise the implanted lifetime of an IC encapsulated in silicone. We did not seek to directly study the degradation of the passivation layer but whether the functional area of the IC (layers below the passivation) was affected by months to years of continuous immersion.

### 2.2. Interdigitated Combs, Surface Conductivity, and EIS Principle and Interpretation

Electrical Impedance Spectroscopy (EIS) can be used to characterise the impedance of electrodes or insulators. The data is often presented as a Bode plot of the complex impedance.

The conventional method for testing the insulation on planar micro-electronic devices is to fabricate an interdigitated comb (IDC) structure in the top conductor layer and then coat it with the protective dielectric insulation. This method has long been used to characterise changes in encapsulation layers in response to varying environmental conditions [13, 52, 55]. When the IDCs are covered not only by passivation layers but also silicone encapsulation, and the samples are immersed in liquid, the impedance between the combs is formed of the layers deposited over them, as well as the encapsulation and electrolyte. The dimensions of the combs affect the sensing depth (i.e. which layer(s) the impedance is most sensitive to) [35, 41]. This principle was applied here to design an IDC to monitor specifically the condition of the dielectric directly above the comb, but not the other layers above that, i.e. the EIS is not sensitive to changes in the insulating layers above the shield, as explained in Section 3.1.

As shown in Figure 2, the Bode plots of the measured impedance are similar to those of a small capacitor, formed by the dielectric between and above the IDC. If liquid water enters, the impedance will decrease as a film forms over the combs, creating a resistive path in parallel with the pre-existing capacitance. Since this process progresses very slowly, we originally imagined an arbitrary threshold for “insulation failure” as a decrease in the magnitude of the impedance by a factor of 10 from the unaged measurement, that is, below 10 MΩ at low frequency (10 mHz). Alternatively, open-circuit failures at the pads are identified as an increase in the magnitude of the impedance.

**Figure 2.**
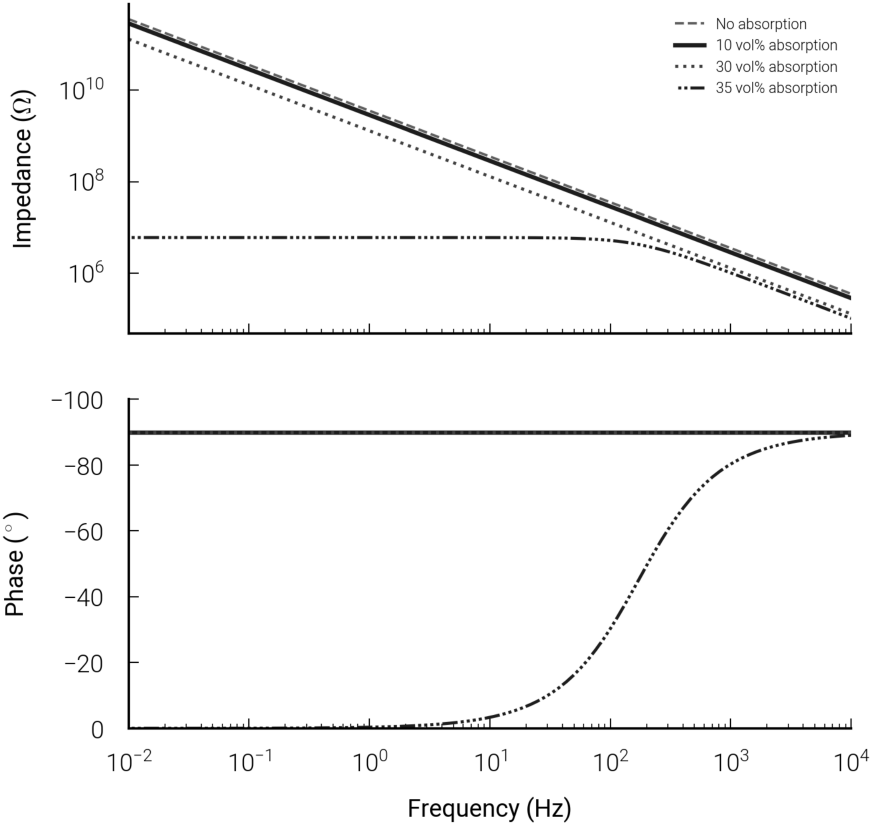
Simulation of the EIS spectrum of a CMOS-IDC at different passivation layer hydration ratios. Reproduced from [40], licensed under CC BY 4.0.

### 2.3. Principle of Accelerated Ageing

Accelerated ageing involves accelerating a dominant failure mechanism by placing samples under elevated stress to gather data, within a reasonable duration, on failures that may take many years to occur under normal use conditions. Caution is needed when planning accelerated ageing experiments, as two competing failure mechanisms may result in the same failure mode. To illustrate, consider one failure mode: open-circuit. It may be caused by two mechanisms: corrosion of a conductive path, or wire bond mechanical “lift off”. In this case, the two mechanisms can be distinguished by visual observation if the IC is not coated with an opaque “glob top”. However, it is not always possible to identify the failure mechanism, nor whether the same mechanism would be dominant in the unstressed situation (as illustrated in Figure 3, and further explained later).

**Figure 3.**
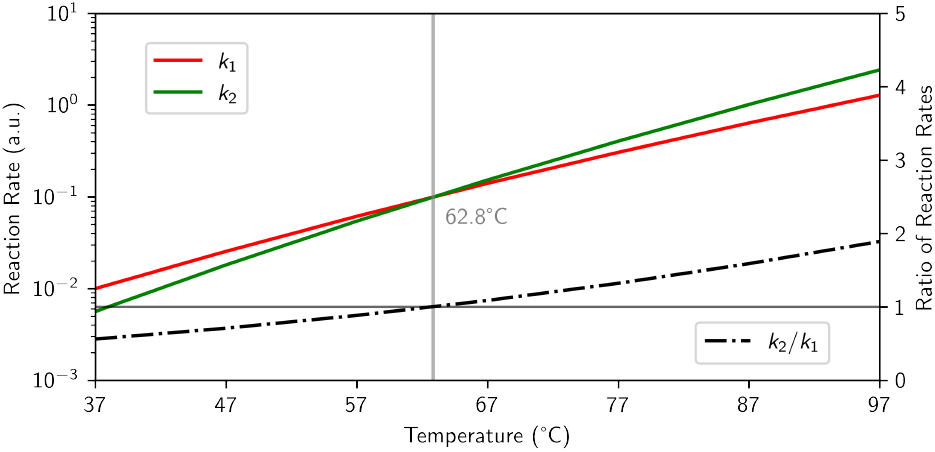
Hypothetical illustration of how, when there are two failure mechanisms (indistinguishable if they lead to the same failure mode), each with a reaction rate *k*_*i*_ ∝ *C*_*i*_ × exp(− *E*_*i*_*/RT*), with different activation energies and constants, the one that dominates at elevated temperature may be slower than the other at working temperature. In this example, *E*_*a*1_ = 0.8 eV, *C*_1_ = 10^11^, and *E*_*a*2_ = 1 eV, *C*_2_ = 10^14^. The dotted line represents the ratio of the two reaction rates. At 62.8^*°*^C, the two rates are equal.

In this paper, we report on experiments where the ageing stress was created by imposing an elevated temperature (47°C, 67°C or 87°C) and a voltage bias (see Section 3.4 for details). Temperature is the most common accelerant, with the assumption that the failure rate *k* of the dominant failure mechanism follows the Arrhenius relationship:

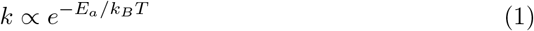

where *E*_*a*_ is the activation energy of the failure mechanism, *k*_*B*_ is the Boltzmann constant, and *T* is the absolute temperature. Extrapolation of the activation energy in operation from tests performed under different conditions requires caution for two reasons:

i. *The activation energy of a failure mechanism is affected by the test conditions*. Tests of IDCs on silicon dioxide have shown that the activation energy varies with the relative humidity [38, 39, 62]. To date there is no universally agreed model of this variability [60]. Our tests are performed on samples immersed in PBS; this condition is representative of the expected working conditions of implanted devices. Reactive oxygen species (ROS) or lipids present in the body may accelerate polymer degradation (see Discussion). However our work suggests that due to the semipermeable membrane properties of silicone rubber, a PBS-only ageing environment is an adequate representation of the implant environment [45]. We do not advocate extrapolating from our data to ICs operating in low relative humidity environments.
ii. *The dominant degradation mechanism can change at higher temperatures*. The acceleration factor used in accelerated ageing experiments must be carefully controlled so that the elevated stress does not alter the dynamics of the failure mechanisms—such as changing which mechanism becomes dominant—or introduce entirely new failure modes. Even in carefully designed experiments targeting a single failure mode, multiple competing mechanisms may coexist. Each mechanism typically has its own reaction rate (assumed here to follow an Arrhenius form):

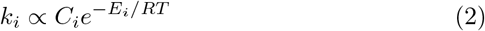

In situations involving competing mechanisms, the failure process with the lower activation energy usually dominates at body temperature. However, at elevated temperatures characteristic of accelerated test conditions, a mechanism with higher activation energy may become dominant. This phenomenon is illustrated in 3, which is intended purely as a general illustration and does not represent data from any specific material. Although specific activation energy data for PDMS is not currently available, similar competing processes have been documented for polyurethane, with a reported change in dominant mechanism occurring around 60°C [4]. Consequently, accelerated tests conducted above this temperature would fail to reveal the primary failure mechanism under operating conditions. Caution must be exercised both when designing accelerated ageing experiments and when interpreting their results. If a transition in the dominant failure process occurs at elevated temperatures, limited insights can be drawn from experiments performed above that transition temperature regarding the likely failure mechanisms at operational temperatures.

The interpolation errors resulting from overly simplified assumptions are illustrated in Table 1. In Scenario 1, an unlikely scenario emerges where a small change in acceleration energy seemingly reverses the expected acceleration trend, suggesting a decrease in lifetime at lower temperatures. This anomaly arises purely from simplifications used in this illustrative example; such behaviour is improbable in practice because real acceleration factors depend on multiple variables beyond temperature alone. Scenario 2 is equally unlikely in practice, as failure mechanisms with lower activation energies tend to dominate at lower temperature. Even when only a single failure mechanism is involved, accurate extrapolation between temperatures is impossible without precise knowledge of its activation energy. Comparing Scenarios 3 and 4 highlights the potential impact of using an incorrect activation energy. If, for instance, the true activation energy is 0.6 eV (Scenario 3), but a higher value of 0.7 eV (Scenario 4) is mistakenly used, perhaps sourced from literature based on different experimental conditions, the predicted acceleration factor, and thus the projected lifetime at 37°C, will be significantly overestimated. In this case, the lifetime would be inflated by approximately 40%, underscoring the importance of context-specific activation energy measurements when interpreting accelerated ageing data.

**Table 1.** Illustration of how different activation energy assumptions affect the true acceleration factor (*A*), compared to the commonly used “rule of thumb” approximation (*A*_*rt*_ = 2^(*T*2*−T*1)*/*10^) with the difference quantified with the error factor: ((*A*_*rt*_ − *A*)*/A*)%. Rows are colour-coded to reflect accuracy: green for close alignment, yellow for mild misalignment, and red for misleading deviation.

| Scenario | $E_{a1}$<br>(eV) | $E_{a2}$<br>(eV) | $T_1$<br>(Body) | $T_2$<br>(Test) | True<br>Acceleration | Rule of<br>Thumb | Error<br>Factor |
| --- | --- | --- | --- | --- | --- | --- | --- |
| 1. Paradoxical Lifetime Reversal | 0.6 | 0.7 | 37°C | 67°C | 0.24 | 8.00 | 3233% |
| 2. Unrealistic Acceleration | 0.7 | 0.6 | 37°C | 67°C | 306.49 | 8.00 | -97% |
| 3. Matched $E_a$, Accurate Estimate | 0.6 | 0.6 | 37°C | 67°C | 7.26 | 8.00 | 10% |
| 4. Matched $E_a$, Mild Underestimate | 0.7 | 0.7 | 37°C | 67°C | 10.10 | 8.00 | -21% |
| 5. Optimal for Rule-of-Thumb | 0.6 | 0.6 | 37°C | 47°C | **2.02** | **2.00** | -1% |
| 6. Mild Overestimation | 0.6 | 0.6 | 37°C | 87°C | 22.63 | 32.00 | 41% |
| 7. High Overestimation | 0.6 | 0.6 | 37°C | 100°C | 44.41 | 78.79 | 77% |

Finally, Table 1 demonstrates the limitations of the commonly cited “rule of thumb,” i.e. “every 10°C increment in temperature doubles the acceleration factor”. As seen in the table, this simplistic approach holds only within a narrow temperature range (Scenario 5). Beyond this range, the rule of thumb significantly overestimates the acceleration factor, potentially leading to overly optimistic predictions of lifetimes at body temperature (Scenario 6).

### 2.4. Failure Modes and Measured Lifetimes

Owing to the ubiquity of microelectronics, the impact of high humidity on integrated circuits reliability has been extensively studied. We previously published a review of the field [60], which we briefly summarise here. A comprehensive list of references is available in our 2013 review; the leading authors of the field are DerMarderosian, Edell, Iannuzzi, Osenbach, Pecht, and Weick.

The definition of device failure varies depending on the specific application. In some cases, failure arises from an open circuit caused directly by corrosion of the metallization layer. In other situations, even a moderate decrease in the insulation properties of the passivation layer can significantly impair integrated circuit (IC) performance, resulting in a different failure mode.

When IC design guidelines are followed, particularly those aimed at minimizing electric field strength across the passivation, corrosion-induced open circuits may be avoided. However, a drop in dielectric insulation resistance alone can still disrupt IC function. This reduced insulation may result from moisture-induced degradation of the passivation layer, which can be accelerated if water-soluble contaminants are present on the passivation surface. When exposed to moisture, these contaminants form a highly conductive ionic fluid, concentrating current flow and accelerating electrochemical degradation.

Specifically, in the presence of moisture, water-soluble contaminants on the passivation surface dissolve to form a conductive ionic solution. This solution then concentrates electric currents at localized points, accelerating electrochemical deterioration of the passivation layer itself.

Applying a conformal coating can improve reliability; however, failures can still occur. Such failures are usually linked either to the formation of a thin liquid film at the interface between the conformal coating and the passivation layer, or to hydration of the passivation material itself. Once initiated, these mechanisms follow pathways analogous to those described for uncoated (bare) ICs.

### 2.5. Previous Experimental Work

We previously explored both custom test structures using non-foundry passivation layers [41] and test ICs sourced from two commercial CMOS foundries [45] to evaluate long-term encapsulation performance.

Lamont et al. [41] investigated the influence of different passivation and encapsulation combinations on insulation impedance. In these studies, samples were subjected to a biphasic electrical bias (± 5 V, corresponding to a field strength of 0.08 V µm^*−*1^) at 67 ^*°*^C. Two groups showed no measurable change in insulation impedance after up to 694 days of exposure: those passivated with either (1) silicon oxynitride (SiO_x_N_y_) or (2) a bilayer of silicon oxynitride and silicon carbide. In contrast, samples passivated with only silicon oxide (SiO_x_) exhibited a significant reduction in insulation impedance over time. An impedance threshold of 10 GΩ was used as the criterion for insulation failure. These electrical observations were corroborated by material analyses including focused ion beam–scanning electron microscopy (FIB-SEM), Fourier-transform infrared spectroscopy (FTIR), and X-ray photoelectron spectroscopy (XPS). It was hypothesised that failure was primarily driven by hydration of the passivation layer via water diffusion through the encapsulant.

In a complementary line of investigation, Nan-bakhsh et al. [45] evaluated the stability of silicone-encapsulated ICs from two CMOS foundries. Custom-designed test ICs were subjected to a year of accelerated in vitro and in vivo testing, with regions of each chip left uncoated to directly assess the inherent hermeticity of the passivated die. Despite continuous electrical biasing in saline environments, the ICs maintained stable electrical performance even in the bare regions, suggesting high intrinsic resilience of modern foundry passivation. Furthermore, it is worth noting that the silicon oxide hydration, which we observed in our in-house IDC devices, was not present in foundry-fabricated CMOS devices suggesting foundry films to be more robust.

However, material characterisation revealed degradation in uncoated die areas, while PDMS-coated regions showed limited deterioration, supporting the viability of PDMS as a long-term encapsulant for ICs in bioelectronic implants. This work highlighted that even permeable elastomers can provide sufficient protection for chronic implantation when combined with robust IC passivation.

Together, these studies provide foundational insights into the failure mechanisms and encapsulation requirements for long-term operation of implantable electronics, forming the basis for the extended investigations presented in this work.

## 3. Methods

### 3.1. Sample Design

#### 3.1.1. IC Technology

CMOS IDCs were fabricated at AMS (Austria Micro Systems AG) in their 0.35 µm CMOS process (C35) with aluminium metallisation and a bilayer passivation of silicon nitride/silicon oxide (see section: “Chemical composition of foundry IC passivation and IMD layers”, page 3 in [45])

#### 3.1.2. Comb Design and Shield Layer

Figure 4 illustrates the horizontal and vertical layouts of the samples. The IDC aluminium metallisation is in the METAL3 layer (typical thickness: 640 nm). Importantly, there is a rectangle of metal in METAL4 (typical thickness: 925 nm), called the Shield, covering the IDC. This was included in the design to protect the active area should the passivation layers become hydrated and degrade. The AMS design rules require regular holes through any metal layer; these are visible in the shield layer in Figure 4A & 4B. The Shield is therefore perforated. Electrically, it is connected to the substrate, as well as to its own bond pad so its potential is determined.

**Figure 4.**
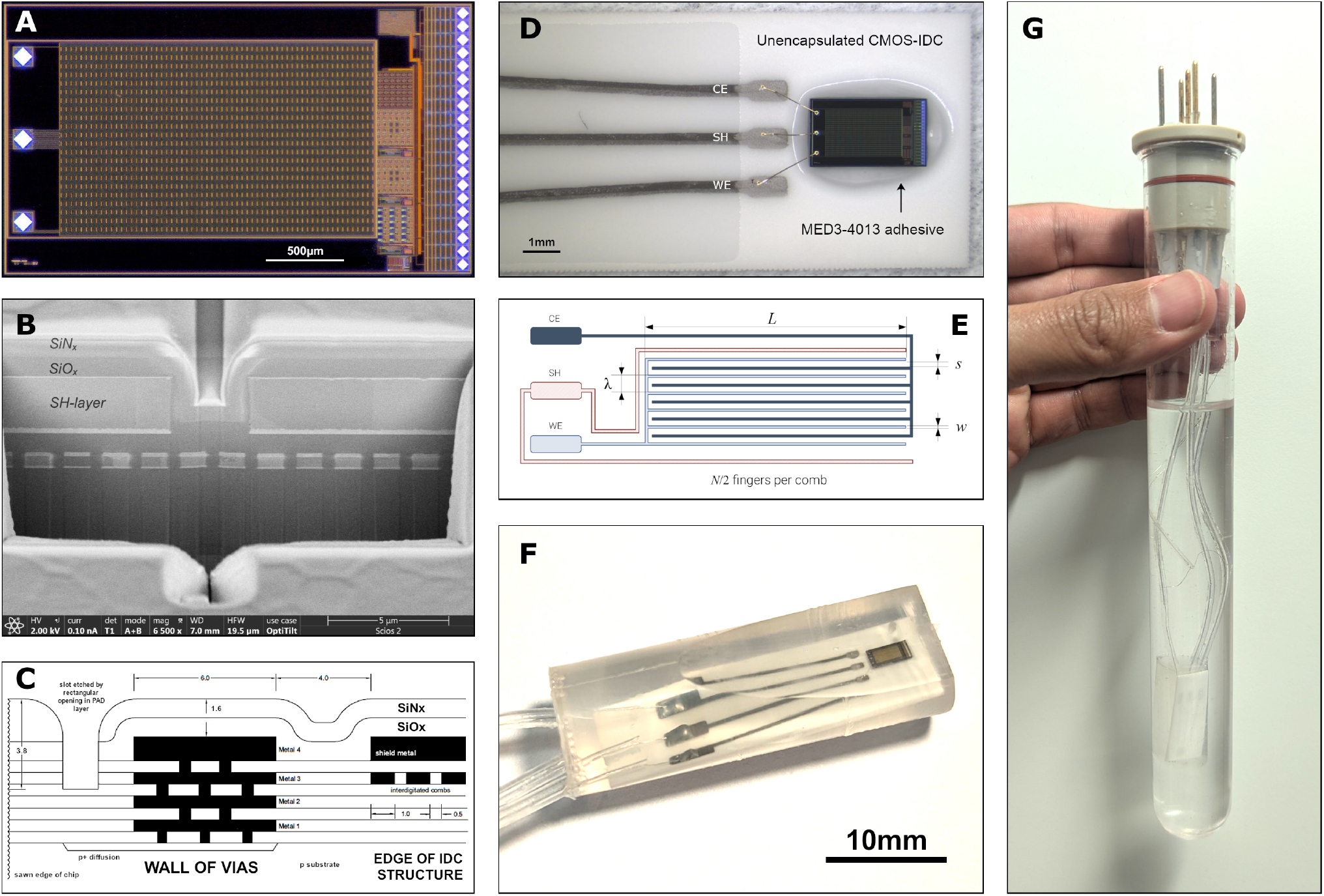
Test IC detail: (A) Optical photograph of test IC, showing the extent of the perforated shield metal, with bond pads to the left, surrounded by a rectangular ‘seal ring’. Outside that, on the right are electrical test structures, and on the outside edge is a second seal ring; (B) SEM image of a FIB cross-section showing regular perforations in the shield (SH) layer of an IC (METAL 4); beneath is the IDC structure (METAL 3). This chip was fabricated using a different process from that used for the samples presented in this study and features a substantially thicker top metal layer (∼ 3 µm vs. 0.93 µm). Included here for illustrative purposes only; (C) Vertical layout of the IC, showing the various metal and dielectric layers, to scale; (D) IC bonded on ceramic adapter with MED-4013 silicone, gold wire bonds to the adaptor pads, and overglaze over the Au/Pt tracks; (E) IDC layout on METAL 3. L = 1.39 mm, s = 0.5 µm, w = 1.0 µm, *λ* = 3.0 µm, N (fingers) = 1450. SH pad and tracks connected to METAL4; (F) Adaptor with IC and wires, encapsulated in MED-6015. Note that this is a photo of an aged sample, the encapsulation has been cut to help visual observation after ageing; and (G) Encapsulated device immersed in PBS in a test tube with three wires connected to the bung (back view, IC not directly visible) Fourth and fifth wires, connecting two depth electrodes to the bung, are also visible.

The IDC, shield and bond pads are surrounded by a rectangular “seal ring”. There is also a second seal ring around the edge of the IC, that encompasses additional test structures (irrelevant to the experiment). The seal ring is a standard structure of rectangular metal and polysilicon tracks at all levels, joined vertically by a dense array of metal vias (Figure 4C). We included this “wall-of-vias” to prevent delamination and lateral water penetration, strengthening the structure by the vertical metal bonding, and forming the connection to the p-substrate through a rectangular p+ diffusion [36]. The final etch which opens windows over the bond pads is applied also around the outside of the seal ring and this penetrates through the passivation layers and some of the inter-layer dielectric (Figure 4C). The wall-of-vias may therefore also prevent cracks propagating from the diced edge into the active area of the chip.

### 3.2. Sample Preparation

Teflon-insulated stainless steel wires (75 µm thick) were soldered (*Hydro-X Multicore* SnPb 60/40) to adaptors made by printing thick film Pt-Au pads and tracks (ESL 5837 G) on ceramic substrates (*Coortek* 96% alumina ADS96R), with overglaze (ESL 4026) over the tracks. The adapters and IDC chips were then cleaned by sonication in a solution of 97 wt% deionised water + 2.5 wt% Na_3_PO_4_ + 0.5 wt% multi-purpose detergent (Teepol™), then acetone (99.8%, Sigma Aldrich), isopropanol (99.5%, Fisher Scientific) and DI water (15 MΩ.cm), for 5 minutes each. The chip was glued on the adaptor with *Nusil* MED-4013 and cured at 150°C. Electrical connection between the two components was achieved with 30 µm diameter gold wire bonds, see Figure 4D (K&S 4524 Ball Bonder, Kulicke & Soffa Ltd, PA, USA).

Samples were then cleaned in an ultrasonic bath as follows: 1 min in acetone, 1 min in IPA, then 1 min in deionised water, then blown dried with nitrogen gas and further dried at 70°C for at least 2 hours.

#### 3.2.1. Silicone Encapsulation

Immediately prior to encapsulation, the samples were held in a *Diener Zepto* Plasma Unit with air plasma for 10 minutes (4 mBar, 100 W) then immediately encapsulated with MED-6015. Part A and B were mixed using a *SpeedMixer* at 2500 rpm for 5 minutes as the plasma surface activation was occurring. The silicone rubber was then cast round the CMOS chip and the ceramic adaptor, including the solder joint to the stainless steel wire, in a cylindrical PTFE mould. A vacuum centrifuge was used to remove bubbles and de-gas [24] before curing at 150°C (room air, atmospheric pressure) for 15 minutes; see Figure 4F.

After encapsulation, the stainless steel wires from the adapters were soldered to the pins in the test tube bung, two depth electrodes were also soldered to two other pins on the bung, and the pins and base of the bungs were encapsulated in *Dow Corning®* 3140 RTV silicone.

After manufacture, every sample was visually inspected and photographed (low magnification) for obvious signs of poor assembly, then each was put in an empty test tube. With the sample dry and at room temperature, EIS data (see Section 3.3) was then collected, from 100 kHz down to 100 mHz to check that the ICs were correctly connected (no short nor open-circuit). PBS (Sigma-Aldrich) was then poured into the tube, immersing the sample (Figure 4G). EIS measurement was done again at room temperature (on immersion day), from 100 kHz down to 100 mHz, then the tube was placed in the heated tank.

### 3.3. EIS Setup

The experiments reported in this paper were performed using the ALTA, a dedicated system that has been described in detail [16].

EIS is performed using a Modulab XM system in a two-electrode setup (see Supplement), applying a sinusoidal voltage between the Counter Electrode (CE) and the Working Electrode (WE) terminals, whilst measuring the current flowing into the WE terminal. There is also a connection from LO to the Shield, to prevent leakage currents not at the IDC from flowing through the femtoammeter [16]. The shield comprises: cable sheaths, tracks on the ALTA modules, a tube around the WE pin at the bung, a track on the adaptor and the metal above and below the IDCs in the CMOS chip.

Figure 5 shows diagrammatically one CMOS IDC sample, connected via its adaptor to the feedthrough pins on the test tube bung. Outside the tube, the multiplexing module is represented with the femtoammeter and the potentiostat. Because the impedance of the femtoammeter is relatively small compared to the insulation being measured, the potential of WE is close to LO, so the impedance being measured is from the CE comb to the WE comb, the shield metal above, and the substrate below. Because the shield metal layer and the substrate are grounded on the chip (see Section 3.1.2) during EIS measurements, the electric field is confined between these layers and will not extend into and beyond the passivation layers. Therefore, the EIS data will not be sensitive to encapsulant delamination or water condensation on the passivation.

**Figure 5.**
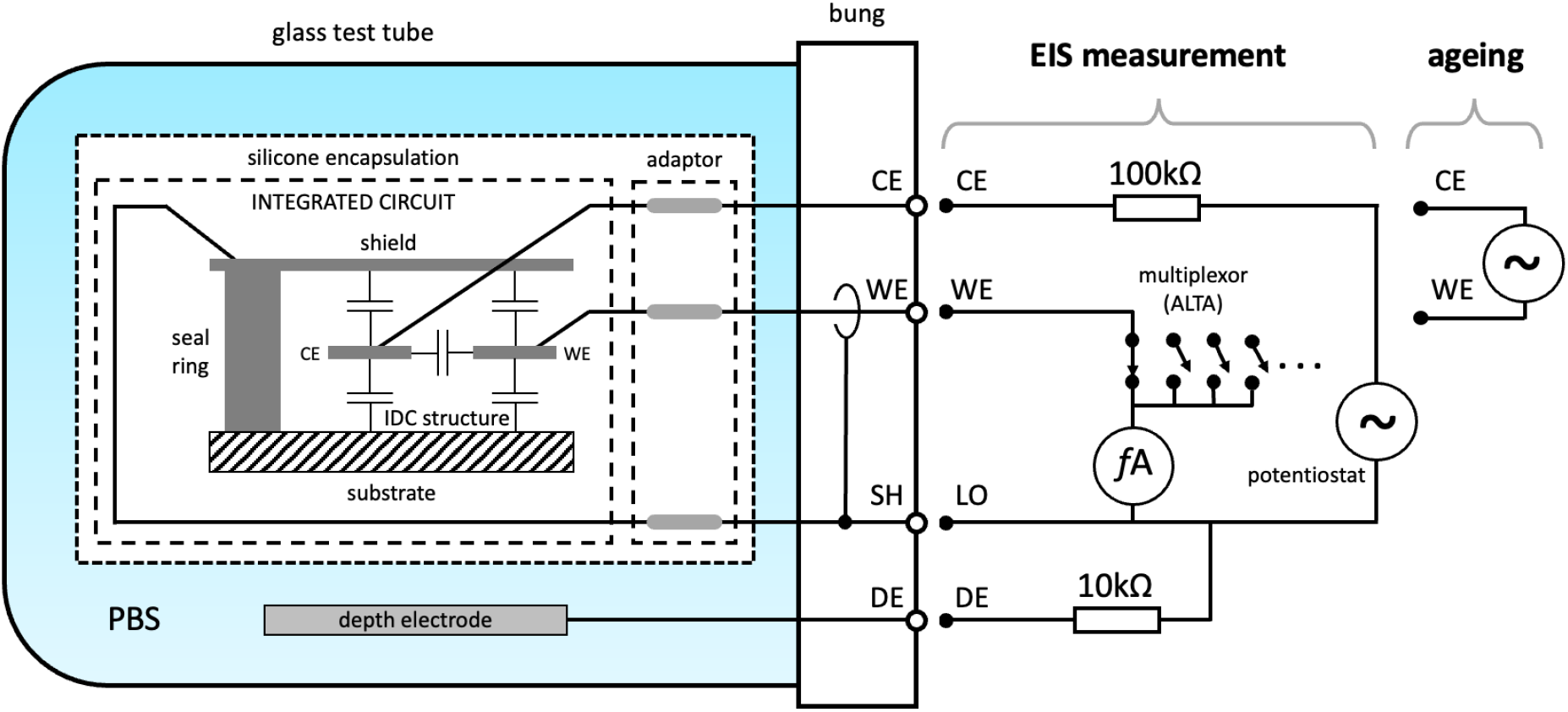
Schematic of one sample: the passivation is represented as capacitances between the substrate, the two combs, and the shield (SH). The three bond pads are marked CE, WE and SH. During impedance measurements, connections are made from the source voltage (potentiostat) to CE and a femtoammeter with WE guarded, as shown. During this measurement, WE and SH are at approximately the same potential. The potential of the PBS is defined by connecting the DE (depth electrode) to SH through a 10 kΩ resistor. Most of the time, the ageing voltage is applied between CE and WE with SH floating. Because of the capacitances of the IDC structure, in this situation, while no deterioration has occurred, the potential at SH is approximately mid-way between CE and WE. When the ageing voltage is DC, positive connection is to CE. See [16] for further apparatus detail.

For the data reported in this paper, the EIS measurements were taken by applying a 50 mV RMS sinewave with frequency sweep from 100 kHz down to 0.01 Hz. This is a field strength of 140 kV/m RMS, applied across the SiO_x_ interlayer dielectric (ILD) between the aluminium combs. There is a similar vertical field strength between METAL3 and METAL4, and between METAL3 and METAL2.

The IDCs have a capacitance of about 150 pF compared to about 6 pF for the ALTA with no IDC connected, as seen in the control samples. Any water that penetrates the sample and bridges the IDCs will appear as a shunt resistance (or perhaps constant phase element) apparent at the low frequency end of the spectrum.

Samples without deterioration should have an EIS spectra characteristic of a ∼ 150 pF capacitor from the lowest frequency (10 mHz) up to about 100 Hz where the magnitude approaches a plateau and the phase changes from -90^*°*^ to 0^*°*^. The plateau resistance would be 100 kΩ if there were only one sample present, but we showed that with several samples connected to the module, the resistance increases proportionally to the number of other samples, giving values close to 1 MΩ (see Eq. 8 in [16]). At even higher frequencies, the RC model exhibits inductive behaviour, with increasing impedance magnitude. This behaviour also depends on the capacitances in other samples in the module.

### 3.4. Experimental Conditions and Batch Descriptions

We performed tests on 4 batches each containing 10 to 12 samples. For continuity between previous publications from the ALTA experiments [41] we used a unique label for each batch. Hence, we refer in this paper to Batches G, H, I, and J. All samples in these batches are made from the same design of CMOS IDC (Section 3.1), prepared as described in Section 3.2. As shown in Figure 4G, the samples were immersed in PBS, each in their own test tube held at three different temperatures: 47°C (Batch I), 67°C (Batches G and H) or 87°C (Batch J). The PBS was prepared in a cleanroom environment from deionised water and PBS tablets (Sigma Aldrich).

Each batch comprised a maximum of 12 samples (see Table 2), a control “sample”, and one empty channel. Each sample is given a unique ID number. Control “samples” are test tubes, filled with PBS and with depth electrodes connected to a bung (with all bung connections encapsulated) but with neither an IDC nor a ceramic adaptor. The bung is connected to a channel of the ALTA, enabling the collection of data to monitor background noise. The empty channels had no connection to the ALTA, providing information on the sources of interference.

**Table 2.** Summary of batches & experimental conditions.

| Condition | G | H | I | J |
| --- | --- | --- | --- | --- |
| PBS Temperature | 67°C | 67°C | 47°C | 87°C |
| Ageing Voltage | ± 5 V | 5 VDC | 5 VDC | 5 VDC |
| Number of Samples | 10 | 11 | 12 | 12 |
| Max Duration (days) | 1286 | 1419 | 1587 | 1582 |

The samples are subjected to an ageing voltage between the CE and WE, i.e. between the two interdigitated combs (see Figure 5), with the shield and depth electrode left floating during ageing [16]. In Batches H, I, and J, the bias was a continuous 5V DC voltage, whilst in Batch G the bias was a 500 Hz biphasic waveform: +5 V, then 0 V, -5 V and back to 0 V, each for 25% of the period. The reason for using the biphasic waveform was that it approximates an optimal drive voltage for LEDs (a legacy from the CANDO project), the negative phase being present to reverse electrochemical processes that may occur. The bias, whether biphasic or DC, was applied continually, interrupted only for the EIS data collection.

An EIS spectrum of each sample in a batch was recorded daily for the first five days, then once a week for the following 3 weeks, then monthly for the duration of the experiment. This ensured that the samples are under bias for the majority of the experiment and also that infant failures may be captured.

When EIS data indicated a failure of the sample, it was taken out of the ALTA, placed (still in the test tube) in a small separate Faraday cage and characterised at room temperature by measuring the EIS between WE and CE as well as EIS between the Shield and a depth electrode. The sample was then dried and photographed, still encapsulated, before any more destructive investigation.

This experiment was affected by the COVID-19 pandemic. During this period, the samples remained immersed in PBS, at room temperature, without any applied bias. No EIS data was collected. Ageing is considered interrupted during this period, i.e. the number of ageing days was not incremented. Batch G was off for 208 days, Batches H and I were off for 209 days and Batch J was off for 210 days.

### 3.5. Statistical Failure Analysis

Life-testing results were analysed using Weibull statistical methods, as they are well established parametric methods for survival analysis, that can handle right-censorship of data. Failures were classified based on their relevance to the failure mechanism studied. Samples removed from the experiment due to causes not related to the failure mechanism under study are “right-censored”: they are counted as surviving up to the day of removal, but their removal is not categorised as a failure. Weibull analysis requires a minimum of two failures. When this condition was met, reliability was quantitatively evaluated using the Weibull distribution:

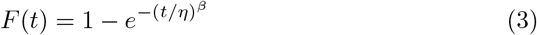

where *η* is the scale parameter (characteristic lifetime) and *β* is the shape parameter (indicating failure rate dynamics). All Weibull analyses were performed using Matlab (release 2022b).

## 4. Results

We tested a total of 45 samples under continuous immersion and electrical bias for durations ranging from 3.5 to 4.3 years (Table 2). Due to the large number of samples and the volume of data collected per sample, individual results are not reported here; a comprehensive table of all results is provided in the Supplement. Instead, we present an overall summary supported by key tables (Tables 3 and 4). This is followed by a focused analysis of stable samples showing no measurable change, as well as control samples (Sections 4.1 and 4.2). We then describe the observed failure modes, including open circuits, dielectric insulation degradation, and pad corrosion.

**Table 3.** Summary of sample faults and failures. These are of four types: (1) open-circuit failures at wire bonds; (2) intermittent open-circuits; (3) delamination at an adaptor leading to CE-SH leakage; and, (4) leakage currents WE-SH at the bung.

| ID | Batch | Fault Day(s) | Retirement Day | Fault/Failure Mode | Comment |
| --- | --- | --- | --- | --- | --- |
| 91 | G | 0 | 0 | Wire bond o/c | Infant failure |
| 103 | H | 71 | 71 |  | o/c failure at CE, no delamination |
| 110 | H | 447 | 447 |  | o/c failure at CE, no delamination |
| 128 | J | 2 | 2 |  | Infant failure |
| 131 | J | 1000–1200 | 1582 | Intermittent o/c | Recovered after Day 1200; no fault at Day 1582 |
| 137 | J | 0 | 500 |  | Recovered after Day 0; no fault at Day 500 |
| 138 | J | 0–7 | 500 |  | Recovered after Day 7; no fault at Day 500 |
| 132 | J | 45 | 45 | Delamination at adaptor | CE-SH ($\sim 1 \text{ M}\Omega$) |
| 106 | H | 71 | 71 | Bung leak | WE-SH leak ($\sim 1 \text{ G}\Omega$); probably at bung |
| 108 | H | 441 | 441 |  | WE-SH leak at bung |
| 136 | J | 1582 | 1582 |  | WE-SH leak at bung |

**Table 4.** Overview of visual observations of corrosion of the samples remaining at the end of the experiment. The results include both absolute counts and percentages of the total number of samples per batch. What constitutes a stable/unstable EIS or a change in visual condition is defined in Sections 4.2 onwards. See Supplement for full description of conditions.

| CATEGORY | CONDITION | G (n=9) | H (n=8) | I (n=12) | J (n=8) |
| --- | --- | --- | --- | --- | --- |
| **Stable EIS**<br>(n=31; 84%) | No corrosion observed | 4 (44%) | – | 6 (50%) | – |
|  | Visible Pb/Sn solder corrosion only | 1 (11%) | – | – | – |
|  | Visible Al bond pad corrosion only | 3 (33%) | 4 (50%) | 4 (33%) | – |
|  | Visible solder & bond pad corrosion | 1 (11%) | 1 (13%) | 2 (17%) | 5 (62%) |
| **Unstable EIS**<br>(n=6; 16%) | No corrosion observed | – | – | – | – |
|  | Visible Pb/Sn solder corrosion only | – | – | – | 3 (38%) |
|  | Visible Al bond pad corrosion only | – | – | – | – |
|  | Visible solder & bond pad corrosion | – | 3 (38%) | – | – |

Table 4 summarises the visual observations made on samples that remained at the end of the study, categorised by whether changes were detected in their electrochemical impedance spectroscopy (EIS) profiles. Notably, the table highlights the absence of a clear correlation between changes in EIS and various visible alterations, which are explored in detail in the subsequent sections.

### 4.1. Stable Samples

Overall, 40% of the samples in Batch G and 50% in Batch I appeared unaffected by the ageing environment. These samples exhibited no changes in either their impedance spectra or visual appearance. This level of stability is particularly striking given their prolonged exposure, 1286 days for Batch G and 1587 days for Batch I, highlighting their exceptional resistance to both electrical degradation and corrosion. Among the remaining samples in these batches, changes were limited to visual alterations only. In Batch G, aside from a single early (infant) failure, no EIS instability was observed. In Batches H and J, 45% and 42% of the samples, respectively, also exhibited only visual changes, while maintaining electrical stability throughout the testing period.

Figure 6 shows a representative stable EIS plot from ID102 (Batch H); both impedance and phase are very stable. Figure 7 shows a pristine ID104 (Batch H) after 1419 days at 67°C under 5V DC bias as a representative of a sample with no visual evidence of degradation: no IC pad corrosion, Pt/Au solder pad corrosion, nor silicone delamination is observed.

**Figure 6.**
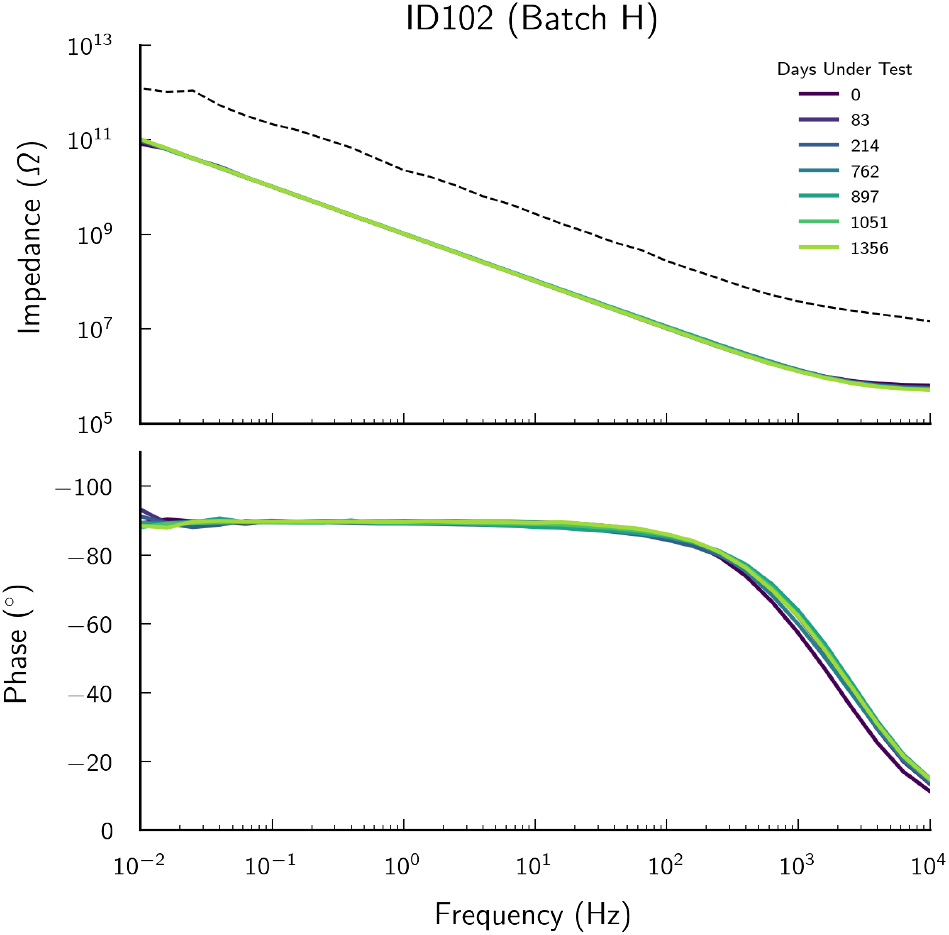
A representative set of stable EIS plots from one sample, ID102 (Batch H), showing a characteristic capacitive response across seven time points; the dashed black line is a data point from a control sample, presented for comparison.

**Figure 7.**
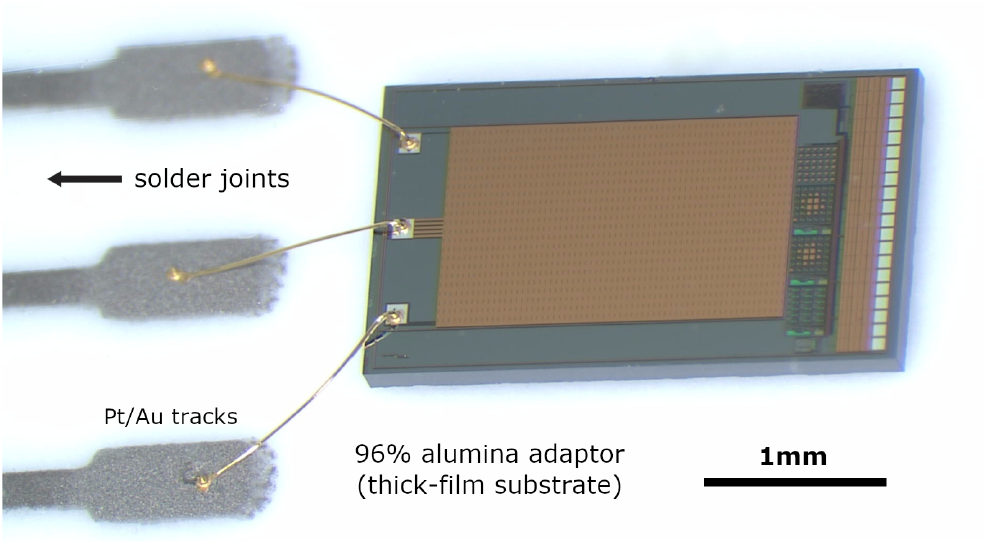
A representative visually pristine sample post-ageing (image taken through transparent MED-6015 silicone): ID104 (Batch H) after 1419 days at 67°C under 5V DC bias, no IC pad corrosion nor Pt/Au pad silicone delamination.

**Figure 8.**
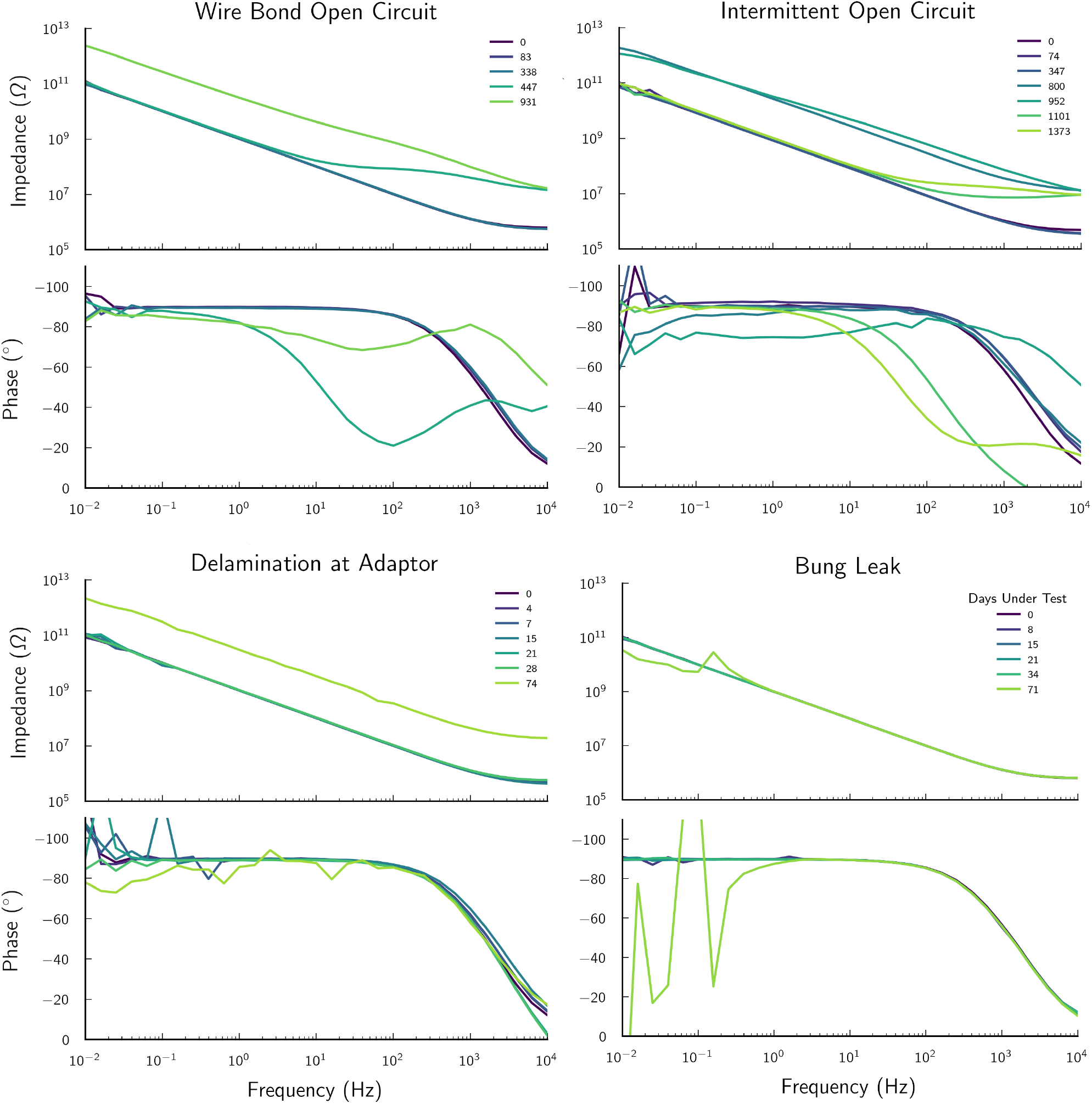
Representative EIS spectra of the four modes of failure/fault observed in the IDC samples. *Top Left:* wire bond open circuit failure in ID110 (Batch H) at Day 447. *Top Right:* intermittent wire bond open circuit failure in ID131 (Batch J). *Bottom Left:* delamination causing a leakage between the CE and SH Pt/Au pads on the ceramic adaptor, with an impedance of ∼ 1 MΩ in ID132 (Batch J); the leakage causes an apparent increase in the impedance (see Supplement). *Bottom Right:* a leak at the bung between SH and WE in ID106 (Batch H) (see Supplement).

### 4.2. Controls

To validate the EIS measurements, a control sample is included in the ALTA for every batch of samples under test. This control consists of an empty tube, prepared in the same way as the sample channels (filled and sealed with a bung), but without a sample: only the two depth electrodes are connected below the bung. For comparison, Figure 6 includes the EIS spectrum of a control sample (ID140, Batch J, test on Day 4) as a black dashed line alongside the data from a stable sample (ID102). The impedance magnitude of a control channel is more than one order of magnitude higher than that of good samples and serves as a benchmark for identifying when a sample has failed as an open-circuit.

### 4.3. Failed Samples

The regular EIS data identified permanent failures or temporary faults (including equipment faults) in 11 of the 45 samples, as summarised in Table 3. Four were open-circuit wire bonds but two of these occurred on Day 0 or Day 2 and we consider these infant failures due to poor bonding rather than due to ageing. We classified three as intermittent open-circuits (all in Batch J). One sample (ID132) showed an impedance of ∼ 1 MΩ between the CE and SH. On visual observation, silicone delamination was observed both at the Pt/Au pads on the ceramic adaptor, where the area delaminated was shorting the CE and SH. Delamination was also observed at the wire-bond pads on the IC, however this did not short the pads and the failure is recorded as delamination at the adaptor. Finally, three samples developed leakages from SH to WE, two definitely at the bung and one (ID106), probably at the bung.

In Section 3.3, we described how failure of the passivation layer by hydration should lead to a decrease in the impedance across the spectrum. We did not observe this in any sample; again we emphasise, we observed no insulation failures in any sample over 3.5 to 4.3 years of ageing.

### 4.4. Reliability Statistics

Table 3 provides an overview of the sample failures and faults observed during the study. For the purpose of studying the underwater reliability of silicon ICs encapsulated in silicone rubber, some of the faults are considered as unrelated to the immersion and failure mechanisms studied, and those samples are right-censored in the reliability analysis. This includes the infant failures at day 0 (Batch G) and day 2 (Batch J) and the three failures attributed to leaks at the bung (Batch H, days 71 and 441, and Batch J, day 1582). The rest are interconnection failures. Since our samples were so stable that we did not observe any insulation failures, we are categorising the interconnection failures as failures (hence the failure mode studied in the survival analysis is no longer insulation failure). All non-failed samples are also entered as right-censored at the completion of the study.

Since there are no failures in Batch I, and no relevant failures in Batch G, Weibull analysis cannot be used to calculate survival data for those batches.

For Batches H and J, the Weibull parameters (*η* is the scale parameter and *β* the shape) were *η* = 51.5 years with *β* = 0.56 in Batch H (67°C) and *η* = 12.5 years with *β* = 0.74 in Batch J (87°C).

### 4.5. Solder and Al Bond Pad Corrosion

We completed a visual inspection of all the samples still under test after 3.5 to 4.3 years (i.e. all the samples except the eight samples that had been removed for further analysis earlier during the experiment). An overview of the outcome of the visual observations is included in Table 4, with details of which of the CE, WE or SH pad was corroded presented in Table 5. The full analysis is in Table S1 in the Supplement.

**Table 5.** Corrosion of Al bonds pads and Pb/Sn solder joints by pad and batch. For bond pads, under microscopic examination, corrosion was categorised into: none (=0), slight (=1), extensive (=2) or total (=3). These were added for each group and sum expressed as a percentage, where 100% would mean all pads were totally corroded, and 0%, no pads corroded. Similarly, for solder joints, pads were sorted into three categories: bright (=0), grey (=1) or black (=2). These were added for each group and sum expressed as a percentage, where 100% would mean all pads were black, and 0%, all pads were bright.

| Batch | Pad | IC Bond Pad | Solder Joint |
| --- | --- | --- | --- |
| G (67°C) | CE | 30% | 11% |
|  | SH | 0% | 6% |
|  | WE | 19% | 6% |
| H (67°C) | CE | 54% | 0% |
|  | SH | 13% | 6% |
|  | WE | 25% | 0% |
| I (47°C) | CE | 19% | 4% |
|  | SH | 19% | 8% |
|  | WE | 28% | 8% |
| J (87°C) | CE | 79% | 63% |
|  | SH | 79% | 75% |
|  | WE | 83% | 38% |

27 samples, representing 40% of Batch G, 72% of H (100% of the samples remaining at the end of the study), 50% in I, and 67% in J (100% of the samples remaining at the end of the study) had visible corrosion of the Al bond pad or the solder joint (see Figure 9). Of these, 21 samples had visual changes and no change in EIS. Comparatively, no samples had changes in EIS without visual changes.

**Figure 9.**
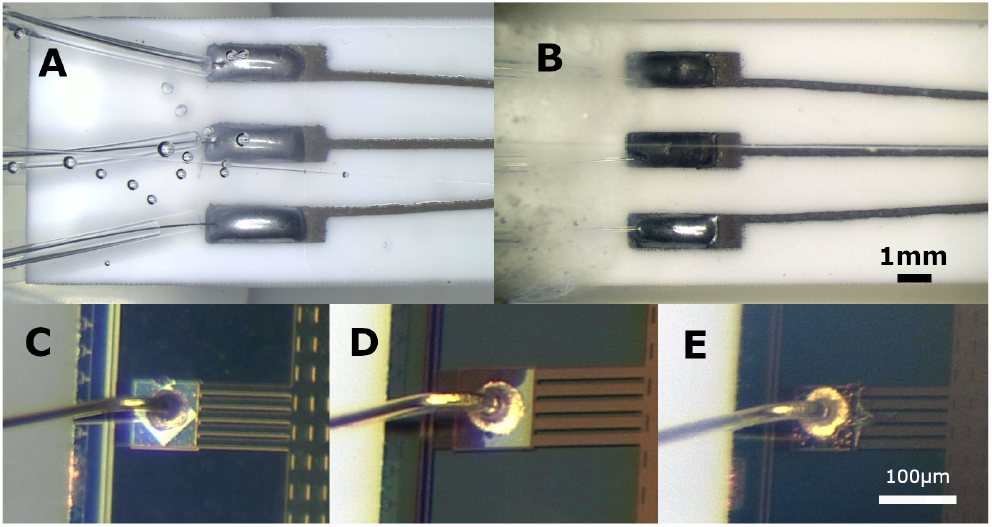
Examples of post-ageing corrosion of solder pads and bond pads; structures within clear silicone. (A) ID086, Batch G, 1286 days, classed as “no visual changes” (note that the bubbles visible are not in contact with any conductive surfaces and hence do not affect the encapsulation efficacy); (B) ID130, Batch J, 1582 days, where the solder appears black at the CE and SH connection, whilst still bright at the WE; (C) ID117, Batch I, 1587 days, SH bond pad is bright, no corrosion; (D) ID118, Batch I, 1587 days, slight SH bond pad corrosion; (E) ID139, Batch J, 1582 days, total SH bond pad corrosion (this did not appear to affect the EIS spectrum).

There were no visible changes to the thick-film pads or tracks of the adaptors. On the adaptors, the tracks are coated with an overglaze layer, only the pads are exposed, hence they may appear lighter on the photographs.

Looking specifically at the Al bond pad corrosion, visible corrosion covered a wider area of the Al bond pads in Batch J samples (87°C). In Batch G (biphasic bias), the corrosion covered less than 50% of the pads except in the only sample that also showed EIS changes (ID093, see Supplement S2). In Batch H (DC bias) all samples with corroded bond pads had corrosion on either CE or WE or both, and three of them also had minor signs of corrosion on the SH pad. A similar trend was observed in Batch I (also DC bias): all samples with corroded bond pads have corrosion on pads CE or WE or both, and three also have signs of corrosion on the SH pad. In Batch J (also DC bias) however, corrosion of the SH pad was present in all samples with corroded pads, and in all of them it was the most corroded pad. In one sample (ID134, with EIS change), corrosion was also observed along the track leading to the SH bond pad.

### 4.6. Changes in the Silicone

The silicone remained optically clear in Batches G (47°C), H (47°C), and I (67°C). In Batch J, immersed at 87°C, there was a yellowing of the silicone rubber in 5 of the 8 samples available for visual observation (Figure 10B).

**Figure 10.**
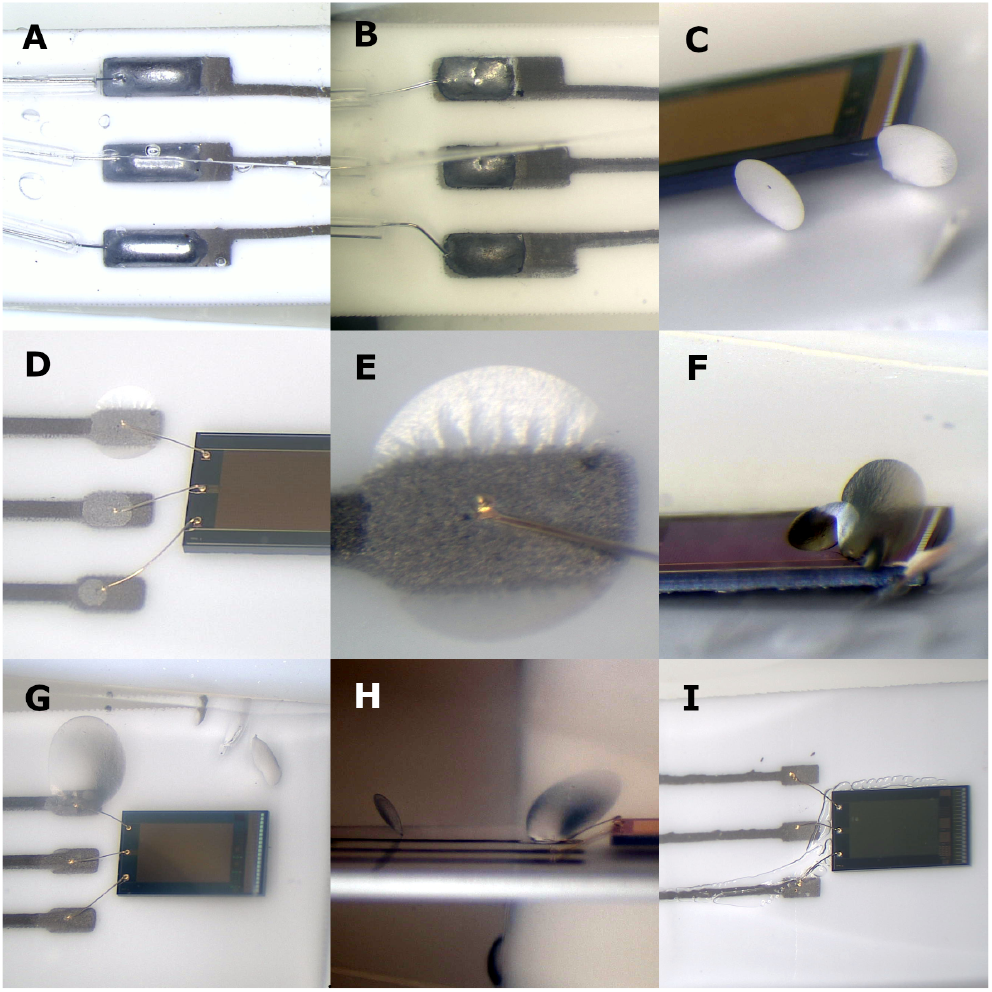
Various rare defects in the encapsulation. (A) clear silicone (ID085, Batch G, 1286 days at 67°C); (B) pronounced silicone yellowing (ID129, Batch J, 1582 days at 87°C); (C) cracks in the silicone bulk (ID129, Batch J); (D) silicone delamination over the Pt/Au bond pads (ID136, Batch J); (E) zoom shot of same sample (ID136) over the CE pad showing “blistering” behaviour; (F) “shear voids” or cracks directly above IDC, at the corner of the chip (ID130, Batch J); (G) crack intersecting with Pt/Au pad delamination (ID139, Batch J); (H) oblique view of same sample (ID139) showing crack; (I) single instance of worm-like void beneath the wire bonds, which was due to poor encapsulation, (ID094, Batch G, 1286 days at 67°C) but the EIS remained unaffected.

Also, only in Batch J, we also observed cracks in the rubber, which we called “shear voids”. These were planar and oblong, usually steeply-inclined to nearby rigid surfaces, but not extending to touch the surface. The silicone rubber appears to be cleaving to relieve stress (see Figure 10C & 10F-H). This was observed on 7 of the 8 surviving Batch J samples, and were accompanied by changes in EIS in samples 129, 130, 131, and 134, where the voids occurred over the IDC area as shown in Figure 10F.

Batch J was also the only batch where we observed signs of delamination of the encapsulant over the thick-film pads with the wire bonds to the IC (Figure 10G). This was observed in five samples: ID130 (WE); ID139 (CE); ID135 (CE and WE); and ID132 & ID136 (CE, WE, and SH). Only two samples showed corresponding changes in EIS: ID130, which also exhibited voids above the IDC surface, and ID132, where delamination extended between pads, resulting in a significant leakage impedance of approximately ∼ 1MΩ. We did not expect much adhesion to the thick-film Pt-Au but leakage currents should be prevented if adhesion remains to the alumina substrate. We do not know why it failed in this sample specifically, but the stress due to thermal expansion was greatest in this batch.

## 5. Discussion

### 5.1. No insulation failures on IDCs

The most obvious failure mode of electronic devices operating in a wet environment is insulation failure as water enters the structure and forms conducting pathways. We tested IDCs immersed in saline held at three temperatures (47°C, 67°C and 87°C) and two voltage biases (5 V DC or ±5 V biphasic) for 3.5 to 4.3 years and we did not detect, in any of the samples, any reduction in their impedance (|*Z*| ≈ 10^11^ Ω at *f* = 10 mHz; see Figure 6); thus there was no significant water ingress. As described in Section 3.1, we designed the test chip with a shield layer (a continuous but perforated metal layer deposited above the IDC passivation layer) and also included the wall-of-vias to reduce lateral penetration, but still, as water will diffuse through the silicone encapsulant, we expected that it would penetrate the chip if there were any crevices, pores or pinholes, larger than water molecules.

The fact that we did not observe any such reduction in impedance means that there are no crevices, or at least, that they have no functional effect even after so many years. The AMS chips we tested, with bilayer silicon oxide/silicon nitride passivation, are effectively flawless. Nevertheless, when exposed to an ionic environment (without encapsulation) the silicon nitride layer does dissolve. Our companion paper reports dissolution rates in PBS of 120 nm/month for ICs from one foundry and 22 nm/month for ICs from another [45]. This shows that the silicone encapsulant prevents dissolution of the passivation.

These observations are supported by earlier work, such as that of Osenbach who tested IDC chips (which we assume he produced in his own lab) encapsulated with a silicone described as “Dow Corning RTV” aged in deionized water at 96°C [46]. He found that a minority (16%) failed after 250 hours but there were then no further failures up to 700 hours (when the experiment was terminated). In a second lot, he intentionally produced voids in the encapsulant by shaking the container to introduce bubbles before application. Consequently, the number of failures rose to 63% after 250 hours, with no additional failures observed up to the end of the experiment (1000 hours). Failure occurred because the passivation layer dissolved where the voids in the rubber were touching the IC surface, exposing the underlying metal. 250 hours would be the time taken to dissolve the passivation layer in Osenbach’s samples.

Given these results, we suggest that there are three reasons why we have not observed electrical insulation failures in our IDCs. The first is the presence of the shield layer, so that if the passivation layers dissolved, the IDC tracks would not immediately be exposed; the aluminium shield and the underlying dielectric would also have to dissolve before the IDC were exposed. The second reason is the remarkable insolubility of modern passivation layers, so that, if there is delamination at the interface, it will take years for the passivation layers to dissolve. Finally, given our specific surface preparation, the MED-6015 silicone rubber encapsulation appears to have excellent adhesion to the passivation layer, with no signs of delamination, even after so many years at elevated temperatures in saline [20, 22]

### 5.2. Silicones

Nusil MED-6015 was selected primarily for its optical transparency. The CANDO project [67], which funded this work, required a transparent and biocompatible micro-LED encapsulant that would not interfere with light transmission essential for optogenetic stimulation of transfected neurons. We previously compared its performance with MED3-4013 [41]. It is an unfilled 2-part liquid silicone rubber that cures via platinum-catalysed hydrosilylation at 150°C. It is not advertised as an adhesive and yet has demonstrated remarkable water-resistant adhesion to the IC surface. The vendor (Polymer Systems Technology Ltd) recommended that we use MED-163 primer to bond to gold, but we doubted that we could apply it as an aerosol repeatably, hence our choice not to use it. For comparison, the WFMA device (described later) used Dow 96-083 (self-priming). The P. E. K. Donaldson work used Dow Corning 3140 and 734 [17–25]. Similarly, the work in our companion paper [45] used room-temperature cured Dow Corning 3140 (in vitro ageing tests) and MED2-4213 (in vivo ageing tests). Much work published in biomedical applications written as “PDMS” implies Sylgard-184, which is not an adhesive [2, 4, 8, 14, 47].

We suspect that the MED-6015 adhesion is a chemical bond to the surface enhanced by our air plasma treatment, but at the present we do not know definitively. We did not use a primer or silane adhesion promoter of any kind (which is likely present mixed in the bulk of Dow 96-083 to give it its “self-priming” nature) and yet adhesion was very robust. Condensation cure silicones (such as Dow Corning 3140 and 734) do not cure by hydrosilylation but perform remarkably as encapsulants (historically, though never in tests as long as in this paper) suggesting the vinyl and hydride functional groups are not key to the adhesion.

These condensation cure silicones present silanol groups that condense into siloxane bonds, the same silanol groups plasma treatments induce on the adherent surface. Perhaps the nature of the siloxane bond itself, which explains the robustness of silicone, may be the reason for the robustness of the encapsulation, which we may discover is an extension of the rubber into the surface itself. This may explain why cohesive failure occurs before adhesive failure in the best silicone bonds. Understanding the mechanisms of adhesion will no doubt lead to even better performance (see Section 5.4.2).

### 5.3. Failures and Faults

Whilst no changes indicative of insulation failure at the IDCs were observed, eleven samples failed or were faulty, as listed in Table 3. Seven were open-circuits: two infant wire bond failures, two open-circuits at the CE connection of uncertain cause (no signs of degradation were visible under the optical microscope), and three intermittent open-circuits. One sample (ID132) delaminated at the adaptor and developed a leakage impedance of about 1MΩ between CE-SH, making the apparent CE-WE impedance abnormally high. Three samples developed leaks at the bung between the WE pin and the surrounding shield tube (see Figure 5C and 5D in [15]).

This section will discuss these various failures in subsections on: apparatus, corrosion & silicone-related phenomena.

#### 5.3.1. Apparatus Fault: Bung Leaks

The impedances measured in this experiment are extremely high compared to the insulation likely to be required in typical implanted medical devices. For example, the leak at the bung on ID106 was 1GΩ. These leaks appeared after periods of 71 to 1582 days in the apparatus, with the test tube containing the sample continuously immersed in water at 67°C or 87°C. In future, it might be possible to find a better adhesive for joining the silicone sleeves that surround the WE wire and pin at the bung, in order to prevent this fault. Certainly, these joints need to be very carefully inspected before using them in the apparatus.

#### 5.3.2. Interconnection Failures

Interconnection failure at wire bonds is a known weakness of implanted devices; we have previously reported on gold wire bond failure on aluminium pads [41,45]. The presence of the silicone encapsulant makes it difficult to check whether the wire bonds are still firmly attached to the bond pad or merely touching. In the absence of any other signs of degradation, it is likely that all open-circuit samples (whether infant or gradual) failed at a wire bond. The infant failures indicate that our bonding process was sub-optimal, which is not surprising as we only bond ICs occasionally. Commercial packaging houses would probably have made more reliable joints.

The silicone encapsulant prevents dissolution of passivation by remaining bonded to the surface, but it does not prevent corrosion of the bond pads or the solder. The visible change in these surfaces is dependent on the temperature (see Figure 9). The fact that this corrosion is occurring on the bond pads probably means that adhesion to the encapsulant is lost and therefore that there will be a film of moisture present at the interface. The welded joint between aluminium and gold forms inter-metallic compounds which are brittle but conductive, and grow as the metal atoms diffuse, perhaps forming *Kirkendall voids* due to the differing densities of the various compounds. At high temperatures, this may cause joint failures (open-circuit) even when dry. The presence of water at the joint surface can cause failure at lower temperatures if there is halogen contamination, which would probably be chloride from the PBS or fluoride from residual foundry etchants in this case [34]. The various intermetallic compounds also have different galvanic potentials, so are likely to corrode where two or more are exposed to the water film. There is likely also to be a water film at the encapsulant-gold interface because adhesion to gold is generally poor, so the water film may well bridge the Al-Au joint, leading to galvanic corrosion with the aluminium being oxidised, the potential difference being about 1.3V [33]. These considerations suggest the need for a better design of joint than the gold ball on the aluminium pad. One possibility is to gold-plate the Al pads so that the weld is Au-Au, avoiding intermetallics [34]. Because more of the surface is now gold, the extent of the liquid film may be greater, but the argument in favour is that it is all gold so will not corrode. However, if a small area of Al remains exposed, perhaps at the edge of the pad, corrosion will be rapid because of the large area for reduction on the gold with high current density at the aluminium. Alternatively, using Al wire on the Al pads would also avoid intermetallics, and improve adhesion (because silicones adhere well to Al_2_O_3_ [23]) but the metal at the pad at the other end of the wire must also be chosen so as not to be vulnerable.

More advanced metallisation stacks are also possible; for example, in the under-bump metallization (UBM) process, which prepares IC pads for solder bumps, a metal stack for aluminium pads can include a titanium-tungsten adhesion layer, followed by a nickel-vanadium barrier layer, finally topped with a copper solder-wettable layer [3].

The Troyk architecture [57] utilises gold wire bonds to establish connections between their ASICs and ceramic adaptors in their wireless floating micro-wire array (WFMA) device [28]. Their device has been implanted in a human volunteer for more than a year as the Intracortical Visual Prosthesis (ICVP) with no reported failures, demonstrating their architecture to be robust [5–7, 30, 37]. Their process involves gold bumping their ASIC pads as a foundry-level process, pre-dicing, prior to gold wire bonding (from personal communication with Troyk; published with permission). Thus, the durability of the interconnections can be attributed to the robustness of the gold–gold bonds. This is visible in Figure 11.

**Figure 11.**
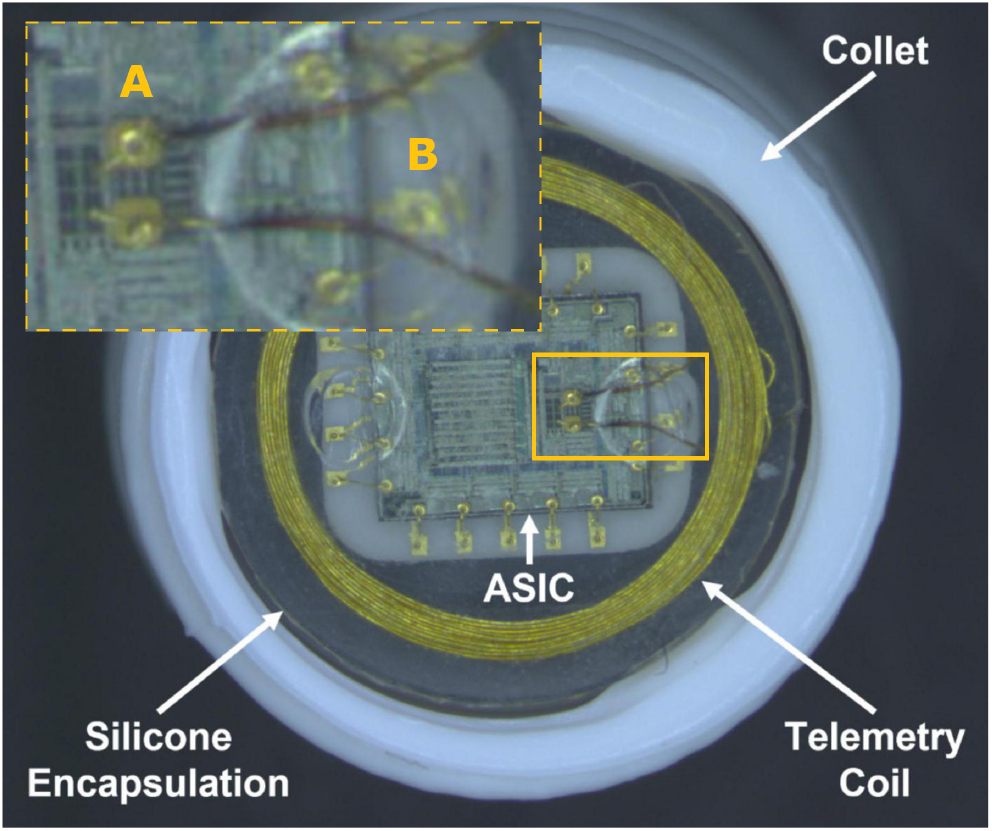
The wireless floating micro-wire array (WFMA) device (zoom emphasis ours): (A) indicating where the chip pads appear metallised with gold; (B) indicating the ceramic baseplate has gold tracks. Reproduced from [28], Frontiers in Neuroscience (2022), under CC BY license.

Thus, we conclude that we might have had fewer conduction failures if the wire bonds had been fabricated by a commercial electronics assembly company and if we had gold-bumped the pads. However, for long-term implantable applications, further research is required to ensure the reliability of encapsulated wire bonds and electronic interconnects operating in wet environments. Comparative studies of various metallisation stacks, wire types, and silicone encapsulants under simulated in vivo conditions are needed. Ideally, the bond welds should remain stable over time, but also strong adhesion should be maintained between the metal surfaces and the encapsulant.

### 5.4. Other Observations

#### 5.4.1. Cracks in the Silicone

This is the first time that cracks in silicone rubber encapsulant have been reported. Although none of them appeared to allow liquid to reach the passivation and cause corrosion, this phenomenon clearly ought to be understood.

The formation of cracks in soft rubbers when placed under triaxial tension was described in 1958 by Gent and Lindley [29]. They found that the stress at which this occurred was proportional to the Young’s Modulus for the rubber. Pourmodheji et al. presented a theory which shows that cracks or spherical voids form under these conditions, depending on the size of pre-existing defects in the rubber, larger defects favouring cracks. Their results show that for a rubber with a Young’s modulus of 10 MPa, which is approximately right, the triaxial stress would need to reach about 50 times atmospheric pressure (5 MPa) to cause cracks, which seems unlikely [50]. However, these papers describe the situation where crack propagation occurs soon after loading whereas in our samples, the cracks may have appeared after a long time in hot water, which may be quite different.

P. E. K. Donaldson compared the time-to-failure of dummy implants made with beam lead transistors with silicone encapsulation in several configurations [18, 25]. He found that the time-to-failure depended on the constraints on the encapsulant: if unconstrained so as to allow changes in volume without large stresses, the devices survived for longer. Thus, thin coatings of ICs should be safe because the encapsulant is not put under stress.

In this experiment, the IDC chips were encapsulated in a mould and once removed, the rubber was free to move perpendicular to the chip surface in order to accommodate volume changes. The mould was made of PTFE, chosen so that the rubber would not adhere to it, but nevertheless, perhaps significant tensile stress was applied as the samples cooled after curing, which led to the cracks immediately or later. However, as shown in Table 6, cracks only appeared in samples from Batch J, strongly suggesting that the high temperature ageing that causes cracking, though whether due to the greater thermal expansion causing greater stress, or due to faster structural changes in the polymer (such as loss of unbound oligomers), we cannot say.

**Table 6.** Number of cracks in the silicone per sample by batch.

| Batch | Bias | n | 0 | 1-5 | >5 |
| --- | --- | --- | --- | --- | --- |
| G (67°C) | ± 5 V | 9 | 9 | 0 | 0 |
| H (67°C) | 5 VDC | 8 | 8 | 0 | 0 |
| I (47°C) | 5 VDC | 12 | 12 | 0 | 0 |
| J (87°C) | 5 VDC | 8 | 1 | 4 | 3 |

**Table 7.** Extent of delamination on adaptor pads by batch: none, slight, extensive, or total.

| Batch | n | None | Slight | Extens. | Total |
| --- | --- | --- | --- | --- | --- |
| G (67°C) | 9 | 6 | 3 | 0 | 0 |
| H (67°C) | 8 | 8 | 0 | 0 | 0 |
| I (47°C) | 12 | 12 | 0 | 0 | 0 |
| J (87°C) | 9 | 4 | 0 | 3 | 2 |

#### 5.4.2. On Adhesion & Blistering

Accelerated aging tests revealed a distinct *blistering* failure mode in ID136 at the counter electrode (CE) pad (see Figure 12) that deserves further attention. The circular morphology of the blister suggests that the delamination is driven by isotropic internal pressure.

**Figure 12.**
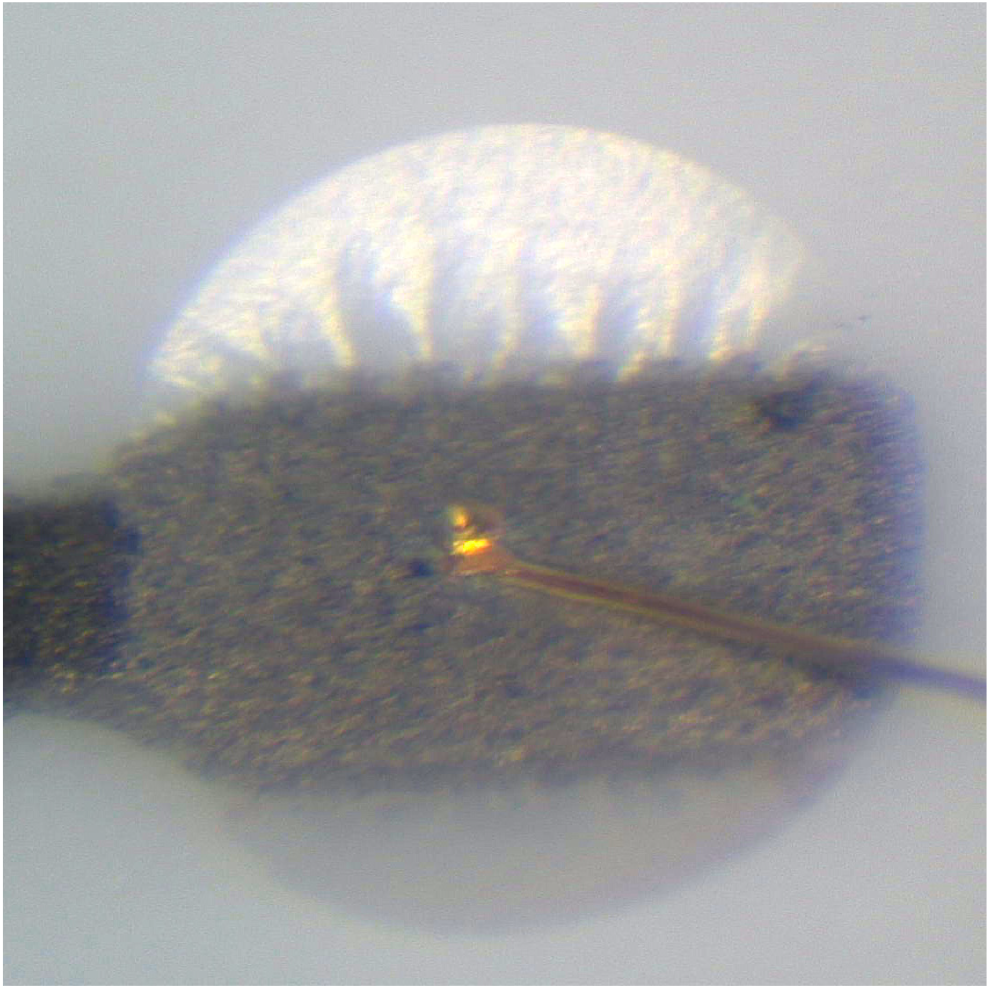
The silicone-encapsulated Pt/Au counter electrode (CE) pad on the alumina adaptor of ID136; blister diameter ∅1 mm.

Similar but less circular failures were observed at pads in other samples aged at 87 °C (ID135, ID139) whereas sample ID130 exhibited delamination exclusively at the working electrode (WE) pad. This selective occurrence exclusively at 87 °C highlights a synergistic effect of elevated temperature and DC bias, required to induce the observed encapsulant delamination.

##### Local Acidification at the Anodic Pad

Delamination was most pronounced at anodic (CE) pads under +5 V DC bias, implicating electrochemical processes. The anodic reaction (2H_2_O → O_2_ + 4H^+^ + 4e^−^) produces protons (H^+^) that locally acidifying the interface, thus weakening silicone-substrate bonds. Oxygen gas evolution additionally induces mechanical stress, creating the blistering effect observed in ID136. Previous work correlated acidic conditions with reduced silicone adhesion under hydrothermal conditions [20] supports this hypothesis.

This was observed over the counter electrode (CE) in ID136 (see Figure 10E). We found that ID136 had failed with a relatively low impedance from WE-SH at the bung. During ageing, SH is usually floating with 5V applied between CE (+) and WE (−). With this bung leak, voltage would appear across CE-SH at the adaptor which would drive electrolysis there at a rate that would depend on delamination between the pads (refer to the circuit diagram Figure 5). Note that blistering of ID136 was therefore probably a double fault, with failures at the bung and on the adaptor, where the blistering occurred.

##### Oxane Bonds and Hydrolytic Equilibrium

A general equation for the condensation reaction of a surface hydroxyl (on some metal M) with a silanol for the formation of an *oxane bond* is:

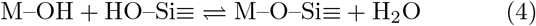

A specific case of this is the bonding of silane coupling agents to silica [49] (or perhaps the passivation of an IC):

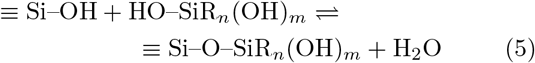

In this equation, *R* refers to a non-hydrolysable organic group, typically containing a functional moiety (e.g. vinyl, amino, or epoxy) that can form covalent bonds with a broader polymer network.

Our theory is that the silicone (MED-6015) adheres to the surface through some process analogous to that of a silane coupling agent reacting with surface hydroxyl groups (which we induced via air plasma). Silane coupling agents anchor polymers to inorganic substrates through the formation of oxane bonds (M–O–Si), yet these linkages are far from immutable. Even fully covalent siloxane (Si–O–Si) bonds are susceptible to hydrolytic cleavage, reverting to silanols with an activation energy of only 98.7 kJ/mol [49]. The propensity for hydrolysis is accentuated by the highly polar nature of the Si–O bond: whereas the Si–C bond possesses roughly 12 % ionic character, Si–O is about 50 % ionic, and M–O–Si bonds to many metals (e.g. Al, Fe, Ti) are still more ionic. In such ionic, inorganic systems the interfacial chemistry is governed less by kinetics than by reversible equilibria; bond formation and breakage proceed according to concentration-driven equilibrium constants.

Water inevitably permeates the polymer/substrate interface, and its impact varies with the surface chemistry of the particular substrate. Silane coupling agents do *not* exclude this moisture; rather, they preserve adhesion by allowing the interface to cycle through hydrolysis and re-condensation. Oxane bonds formed with silica or glass, for instance, hydrolyse during prolonged immersion but re-form upon drying, evidencing a reversible, quasi-equilibrium condition at the interface. Direct proof of true equilibrium is elusive, yet the collective observations of silanol–siloxane interconversions strongly imply that interfacial bonding is dynamic and concentration-controlled rather than permanently fixed [49].

##### Stable Adhesion

Plueddemann outlined three key conditions for favourable siloxane bonding equilibria, i.e., to favour condensation over hydrolysis: (1) maximal initial formation of M–O–Si bonds, (2) a minimal equilibrium concentration of water at the interface, and (3) polymer structures that retain silanol groups near the surface [49]. In our system, condition (1) is addressed by air plasma treatment, which generates a high density of surface silanols, providing abundant potential bonding sites. For condition (2), we implement a drying step prior to encapsulation and cure the samples at 150°C. Furthermore, our high levels of surface cleanliness prior to encapsulation, coupled with the saline ageing environment, ionic concentration gradients are likely to drive osmotic water transport away from the interface, helping maintain low interfacial water levels [19, 26]. Regarding condition (3), our use of vacuum centrifugation during polymer application [24], along with the rigid, static nature of the samples, promotes an intimate and stable contact between the polymer and the substrate. Together, these factors may help explain the exceptionally stable adhesion observed in our samples.

##### Thermochemical Considerations

Preferential failure (delamination) at alumina substrates rather than at the IC may arise from thermochemical stability differences. Bond dissociation energies of Si–O (799 kJ/mol) significantly exceed Al–O (511 kJ/mol) [42], implying greater resistance of silicon-based interfaces to hydrolysis and thermal stresses, perhaps explaining why delamination over the IC is observed in only one sample. As these are strongly exothermic bonds, elevated temperatures further bias equilibrium toward bond breakage (Le Chatelier’s principle) as evidenced by this occurring only in 87 °C samples, and not at lower temperatures.

##### Accelerated Aging vs. Physiological Conditions

The described failure mode, occurring exclusively under combined high temperature (87 °C) and DC bias, exemplifies the potential pitfalls of excessively aggressive accelerated aging protocols. At physiological temperature (37 °C), such failures are unlikely due to slower kinetics of hydrolysis and diffusion-mediated pH stabilization in biological environments. These findings underscore careful selection of accelerated aging conditions to avoid artificially induced failure modes absent at physiological conditions (See Figure 2 & Table 1, main text).

#### 5.4.3. Visible Corrosion on Solder and Pads

The severity of corrosion was strongly associated with temperature, with higher ageing temperatures producing more extensive degradation across both aluminium bond pads and Pb/Sn solder joints (Table 5). The most pronounced corrosion was observed in Batch J (87 °C), where bond pad corrosion ranged from 79–83% and solder joint corrosion from 38–75%. In contrast, moderate corrosion was seen in Batches G and H (67 °C), while the lowest levels occurred in Batch I (47 °C), demonstrating thermal acceleration of a material degradation process.

The extent of corrosion also varied by pad type. Counter electrode (CE) pads occasionally showed higher corrosion than shield (SH) or working electrode (WE) pads under certain ageing conditions. SH pads tended to exhibit lower solder joint degradation, suggesting that electrode position or function may influence localized corrosion behaviour. However, when comparing total corroded pads across all batches, no single bond pad type (CE, WE, or SH) was consistently more affected. This indicates that corrosion was likely driven by local galvanic interactions and material interfaces rather than the direction of current flow between electrodes. Across all but the highest temperature condition, bond pad corrosion exceeded that of solder joints, but this difference diminished at 87 °C, where both interfaces were severely affected— suggesting that multiple degradation mechanisms converge under extreme conditions. Solder joint corrosion was more variable overall, likely reflecting localised influences such as flux residues, alloy composition, or intermetallic growth in addition to thermal and electrical stresses.

Interestingly, not all samples exhibiting visual corrosion at the bond pads showed corresponding changes in EIS data. In fact, 40% of samples in Batches G, H, and I presented visible corrosion without any detectable EIS changes. Conversely, no instances of EIS changes were observed without accompanying visual corrosion. This suggests that solder and/or pad corrosion may precede detectable changes in the EIS spectra. However, since visual inspections were conducted at a single time point, we cannot determine whether EIS variations correlate with corrosion onset over time, nor can we establish whether corrosion initiates preferentially at specific bond pads.

#### 5.5. Survival Analysis

Although no IC insulation failures were observed in any of the batches, several samples were removed from the ageing experiment owing to interconnection failures and equipment faults (Table 3). Using this data, a survival analysis was performed on the results of Batches H (67°C) and J (87°C). Batch G (67°C, biphasic) and Batch I (47°C) were excluded as there were no relevant failures. The shape parameter *β* in both batches is less than 1, indicating that the failure rate decreases with time. This is not surprising given the limited number of failures in each batch, and is unlikely to be true in practice, which calls for care in interpreting the scale parameters (*η* = 51.5 years in Batch H and *η* = 12.5 years in Batch J). Often called the characteristic life, *η* indicates the time at which there is a probability that 62.3% of the samples have failed. In fields where the failure of a device can have catastrophic consequences or where replacement is not an option, it may be more interesting to evaluate the time to reach lower cumulative failure probabilities, as done in Figure 13. These results were obtained under accelerated ageing conditions (H at 67°C, J at 87 °C, both with 5V DC bias). To infer a lifetime under operating conditions, the acceleration factor between the accelerated and operating conditions must be known. It can be derived from survival data under three accelerated conditions (three data points are the minimum to verify the proportional relation between these conditions). Whilst we did implement three different temperatures with the same bias in Batches H to J, there were no failures in Batch I; hence, we only have two data points. Although our limited data does not enable us to verify it, we make the assumption that the failure reaction rate follows an Arrhenius reaction, and thus the survival time is proportional to 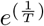. This assumption was used to extrapolate the cumulative wire bond failure probabilities to 37°C in Figure 13.

**Figure 13.**
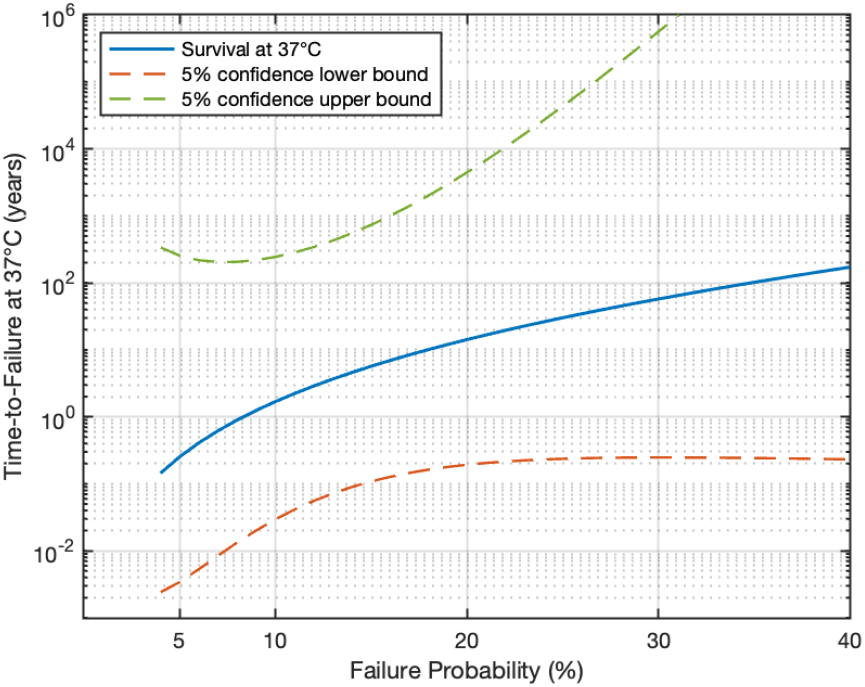
Cumulative probability of wire bond failure at 37°C, interpolated from Weibull analysis of failures at 67°C and 87°C. The large confidence interval can be explained by the low *β* at 67°C.

#### 5.6. The Impact of Accelerating Conditions

Whilst there were no passivation failures, we can draw a few conclusions on the impact of the experimental conditions and likely causes of failure of future silicone-encapsulated IC implants.

Our work suggests that failure of implants designed with ICs passivated with a SiO_*x*_/SiN_*y*_ bilayer and encapsulated with silicone rubber is most likely to be caused by interconnection failure than corrosion of the active area.

Referring to Table 4, temperature does accelerate deterioration. At 47°C, 67°C and 87°C respectively, 50%, 100% and 100% of the samples observed had visible signs of deterioration, but EIS had changed in only 0%, 29% and 38% of the samples.

In the two batches with failures during ageing, H (67°C) and J (87°C), a continuous 5 V bias was applied between the two interdigitated combs. Comparatively, there were no failures in Batch G (67°C, biphasic bias). Likewise, at the same temperature, there were more changes in the continuous bias (Batch H) than in the biphasic bias (Batch G), 29% versus 0% respectively in EIS changes, and similarly for visual observations, the increase is to 100% from 56%. This may be due to the alternating nature of the biphasic bias, or to the 50% duty cycle. We cannot conclude whether it is the duty cycle or the reversal of potential (that may lead to some reversal of the reactions taking place at the interconnections) that contributes to the lower number of changes in Batch G compared to Batch H.

#### 5.7. Study Limitations

The bathing solution used in this experiment was phosphate-buffered saline which matches the pH and osmolality of body fluid but does not imitate all the components of body fluid, and in particular, does not include hydrophobic constituents.

It is known that silicones are impermeable to salts and oxides, while being permeable to nonpolar molecules including solvents (e.g. n-heptane, toluene) [15]. The impermeability to salts means that the encapsulant behaves as a semi-permeable membrane [19]. However, it is also known that silicones can absorb lipids from the body. Carmen and Kahn found that silicone rubber balls exposed to blood in heart valves gained weight at 0.27% per month, reaching increases as high as 41% and sometimes causing them to split [12]. Alterations in mechanical properties have also been reported [56, 61]. There are many hydrophobic compounds in the body that will be preferentially absorbed into silicones rather than water [66].

For implants in soft tissue, the bathing liquid would be interstitial fluid. Fatty acids are transported from the capillary membranes to the cells bound to the protein albumin [58]. The concentration of these fatty acids depends on recent products of digestion. Despite this binding to albumin, the fatty acids are efficiently transferred to cells.

Several questions arise about silicone implants in soft tissue. How much fatty acid will be absorbed by the encapsulant, how will it be distributed, and how quickly will this happen? What will the effect of the absorption be on the bulk properties (modulus, strength, hardness) and on the adhesion to the underlying surfaces (passivation layer, pads, etc.)?

These questions need to be answered before one could be confident about the longevity of encapsulated implants, though no obvious deleterious effects from hydrophobic constituents have been reported in existing implants with silicone rubber encapsulation as far as we know [10].

Another species that might affect lifetime is reactive oxygen, produced by the foreign body reaction. Caldwell et al. [11] have shown that the Parylene-C insulation on Utah Arrays is attacked by reactive oxygen. Although polysiloxanes seem less likely to be affected by a strong oxidising agent, there remains a possibility that it, or other species present in the tissue will have some harmful effect on the device. Life-testing, possibly accelerated, is a most convincing way to show that a device will survive long enough in operation, but if it takes around 5 years to do so, the chip technology is likely to have changed by the time the evidence has been collected. This may mean that implanted devices must be made from IC technology that is not up-to-date. However, the situation may be less difficult if the properties of the passivation, metallisation, encapsulant are known, and certain design features, such as the seal ring and maximum field-strengths, are maintained.

#### 5.8. Designing Implants Based on these Results

Our IDC chips had two features intended to prevent failure when tested in implant conditions: an additional wall-of-vias and a shield. The area needed for the wall-of-vias is small (Figure 4C), may be required by the foundry, and, as none of these IDC chips showed failure by edge-penetration of water, may have been effective. Walls-of-vias should be used in future implant designs.

The advantage conferred by the shield is less obvious because we saw no dissolution of the passivation layer under the silicone and the shield was intended to prolong lifetime if dissolution occurred. Designers must decide whether or not they can implement the circuit without the top metal layer, so it can be used as extra protection.

As we only had failures related to interconnection, one might assume a silicone-encapsulated fully wireless system with power and communication antenna a trace under passivation, such as in Figure 6 of [1], would be an implant architecture that would last long beyond the predictions of this paper.

## 6. Conclusion

This study set out to evaluate the reliability of silicone-encapsulated CMOS ICs under simulated implant conditions, with the expectation that insulation failures due to water ingress would rapidly occur, due to silicone rubber’s high water permeability. Surprisingly, no insulation failures were detected over 4.3 years of accelerated ageing at 47°C, 67°C, and 87°C under electrical bias. Electrical impedance spectroscopy (EIS) revealed no significant dielectric insulation degradation, and the foundry passivation, encapsulated in adhesive silicone rubber, remained intact under prolonged immersion in hot phosphate-buffered saline.

Instead, the primary failure modes were related to interconnection; gold wire bonds on aluminium pads emerged as the most vulnerable component, exhibiting both infant failures and gradual degradation observed as open circuits. Corrosion of some of these aluminium bond pads was observed, and also corrosion of Pb/Sn solder pads; neither were correlated with EIS changes.

These findings provide important insights for the design of polymer-packaged miniaturised implantable electronics. While silicone encapsulation demonstrates strong long-term stability, future implant designs must account for interconnection reliability, and the effect of field strength in driving local electrolysis. Wire bond failures, pad corrosion, and DC-driven electrolysis-induced delamination must be mitigated through careful material selection, packaging strategies, and choices of system architecture.

Two aspects of silicone encapsulation need investigation: the strange cracking seen in the 87 °C samples; and the effects of hydrophobic constituents of interstitial fluid. If the cracking phenomenon only occurs at high temperatures but not at body temperature, it may limit the elevated temperature that can be used for life-test acceleration.

While we are cautious not to extrapolate a specific lifetime at 37°C from these data, the absence of insulation failures after 4.3 years of continuous accelerated ageing under electrical bias and elevated temperatures strongly suggests that silicone-encapsulated ICs, thoroughly cleaned and plasma treated pre-encapsulation, can offer robust long-term performance. These findings provide compelling evidence that, when combined with stable foundry passivation, silicone encapsulation is a credible alternative to traditional hermetic enclosures for many miniaturised implantable applications.

## 7. Acknowledgements

This work was funded by the Wellcome Trust (grant ref: 102037) and the EPSRC (grant ref: NS/A000026/1) as part of the CANDO Innovative Engineering for Health project. It also received support from: Project POSITION-II, funded by the Electronic Components and Systems for European Leadership Joint Undertaking (ECSEL JU) in collaboration with the European Union’s H2020 Framework Program (H2020/2014-2020) and National Authorities (grant agreement Ecsel-783132-Position-II-2017-IA); from Toshiba Research UK who supported CL’s doctoral work (Studentship Number 000026177); from the Wellcome Trust (WT 218286/Z/19/Z); and also from the General Sir John Monash Foundation who funded ASI’s doctoral studentship.

