## Supplement for "On the Stability of Silicone-Encapsulated CMOS ICs for Active Implantable Devices: 4.3 Years of Accelerated Life Testing"

### Supplementary Material

**Abstract.** This document contains supplementary material in support of the paper: "On the Stability of Silicone-Encapsulated CMOS ICs for Active Implantable Devices: 4.3 Years of Accelerated Life Testing".

Supplement 1 presents data from electrical and visual observations from all the CMOS IDC samples. Supplement 2 describes in detail the experimental setup and impedance spectroscopy analysis using a Modulab XM system for repeatability, including an examination of aliasing and low-frequency electrical noise with representative time-domain and Bode plot data.

#### Supplement 1. Sample Data

Table S1 summarises and explains the terms used in Table S2 and S3. Table S2 presents all the visual changes observed in the IDC samples under microscope examination, focussing on corrosion (both of the Pb/Sn solder joints of the alumina adaptor and of the Al bond pads of the IC) and on a variety of silicone changes. Table S3 is a summary of all the electrical changes observed in the EIS spectra of the IDC samples measured through the Solartron potentiostat.

**Table S1.** *Qualitative categorisation criteria for visual inspection and EIS analysis.*

| Categorisation | Terms Used | Explanation |
|---|---|---|
| **EIS Instability** | stable / unstable | EIS is considered unstable if the impedance magnitude $\|Z\|$ varies by $\pm0.5$ orders of magnitude at $f = 10\,\text{mHz}$ across repeated measurements. |
| **Solder appearance** | bright / grey / black | Assesses the visual appearance of the solder joints: "bright" indicates retaining a shine comparable to a freshly formed joint, "grey" indicates a mildly tarnished surface, and "black" means heavily oxidised. |

| Categorisation | Terms Used | Explanation |
|---|---|---|
| **Aluminium discolouration** | none / slight / extensive / total / total+ | The aluminium bond pads of the IC exhibit progressive corrosion, visible as a black, irregular surface. "None" indicates a fully shiny, flat pad as received from the foundry; "slight" indicates $< 50\%$ of the pad area is corroded; "extensive" indicates $> 50\%$ is corroded; "total" means the entire pad area is affected; and "total+" indicates discolouration extends beyond the pad into metallisation beneath the passivation layer. |
| **Silicone colour** | clear / cloudy / yellowing | Assesses the visual appearance of the bulk silicone encapsulation: "clear" indicates transparent silicone with no visual degradation; "cloudy" indicates general loss of optical clarity; and "yellowing" indicates widespread discolouration and increased opacity throughout the encapsulant. |
| **Pt–Au delamination** | none / slight / extensive / total | Assesses delamination of silicone over the Pt/Au pads on the ceramic adaptor where gold wire bonds land: "none" indicates full adhesion; "slight" means $< 50\%$ of pads are affected; "extensive" means $> 50\%$ show delamination; and "total" means all pads are delaminated. |
| **Shear failures (#)** | none / 0–5 / >5 | Refers to the number of shear void failures observed in the silicone: "none" indicates none observed; "0–5" denotes a low count; "$> 5$" indicates frequent or widespread occurrences. |
| **Voids touching IC** | none / yes | Refers to whether any of the shear failure voids are touching the IC. |

| SAMPLE INFORMATION | | | | | | | VISUAL DATA | | | | | | | | | |
|---|---|---|---|---|---|---|---|---|---|---|---|---|---|---|---|---|
| | | | | | | | CORROSION | | | | | | SILICONE CHANGES | | | |
| ID | Batch | Voltage | Temp. | Retirement Day | Fault/Failure Day | Failure/Fault | Solder Joints (Colour) | | | IC Bond Pad (Area) | | | Silicone Appearance | Pt/Au Delamination | Shear Failures (#) | Voids Touching IC? |
| | | | | | | | CE | SH | WE | CE | SH | WE | | | | |
| 85 | G | ±5V | 67°C | 1286 | – | – | bright | bright | bright | none | none | none | clear | none | none | none |
| 86 | G | ±5V | 67°C | 1286 | – | – | bright | bright | bright | slight | none | none | clear | none | none | none |
| 87 | G | ±5V | 67°C | 1286 | – | – | bright | bright | bright | none | none | none | clear | none | none | none |
| 88 | G | ±5V | 67°C | 1286 | – | – | bright | bright | bright | none | none | none | clear | none | none | none |
| 89 | G | ±5V | 67°C | 1286 | – | – | bright | bright | bright | extensive | none | extensive | clear | slight | none | none |
| 90 | G | ±5V | 67°C | 1286 | – | – | bright | bright | bright | extensive | none | none | clear | slight | none | none |
| 91 | G | ±5V | 67°C | 0 | 0 | wire bond o/c | sample removed at Day 0; unavailable for visual inspection | | | | | | | | | |
| 93 | G | ±5V | 67°C | 1286 | – | – | grey | bright | bright | total | none | total | clear | slight | none | none |
| 94 | G | ±5V | 67°C | 1286 | – | – | grey | grey | grey | none | none | none | clear | none | none | none |
| 95 | G | ±5V | 67°C | 1286 | – | – | bright | bright | bright | none | none | none | clear | none | none | none |
| 99 | H | 5VDC | 67°C | 1419 | – | – | bright | bright | bright | extensive | none | none | clear | none | none | none |
| 100 | H | 5VDC | 67°C | 1419 | – | – | bright | bright | bright | total | slight | none | clear | none | none | none |
| 101 | H | 5VDC | 67°C | 1419 | – | – | bright | grey | bright | total | slight | slight | clear | none | none | none |
| 102 | H | 5VDC | 67°C | 1419 | – | – | bright | bright | bright | total | slight | extensive | clear | none | none | none |
| 103 | H | 5VDC | 67°C | 71 | 71 | wire bond o/c | sample removed; unavailable for visual inspection | | | | | | | | | |
| 104 | H | 5VDC | 67°C | 1419 | – | – | bright | bright | bright | slight | none | none | clear | none | none | none |
| 106 | H | 5VDC | 67°C | 71 | 71 | bung leak | sample removed at Day 71; unavailable for visual inspection | | | | | | | | | |
| 107 | H | 5VDC | 67°C | 1419 | – | – | bright | bright | bright | none | none | slight | clear | none | none | none |
| 108 | H | 5VDC | 67°C | 441 | 441 | bung leak | bright | bright | bright | none | none | extensive | clear | none | none | none |
| 109 | H | 5VDC | 67°C | 1419 | – | – | bright | bright | bright | slight | none | none | clear | none | none | none |
| 110 | H | 5VDC | 67°C | 447 | 447 | wire bond o/c | sample removed; unavailable for visual inspection | | | | | | | | | |
| 113 | I | 5VDC | 47°C | 1587 | – | – | bright | bright | bright | none | none | none | clear | none | none | none |
| 114 | I | 5VDC | 47°C | 1587 | – | – | bright | bright | bright | none | none | slight | clear | none | none | none |
| 115 | I | 5VDC | 47°C | 1587 | – | – | bright | bright | bright | none | none | none | clear | none | none | none |
| 116 | I | 5VDC | 47°C | 1587 | – | – | bright | bright | bright | total | extensive | total | clear | none | none | none |
| 117 | I | 5VDC | 47°C | 1587 | – | – | bright | bright | bright | none | none | none | clear | none | none | none |
| 118 | I | 5VDC | 47°C | 1587 | – | – | bright | grey | grey | extensive | extensive | slight | clear | none | none | none |
| 120 | I | 5VDC | 47°C | 1587 | – | – | bright | bright | bright | slight | none | slight | clear | none | none | none |
| 121 | I | 5VDC | 47°C | 1587 | – | – | bright | bright | bright | none | none | none | clear | none | none | none |
| 122 | I | 5VDC | 47°C | 1587 | – | – | bright | bright | bright | none | none | none | clear | none | none | none |
| 123 | I | 5VDC | 47°C | 1587 | – | – | bright | bright | bright | none | none | none | clear | none | none | none |
| 124 | I | 5VDC | 47°C | 1587 | – | – | grey | grey | grey | none | none | slight | clear | none | none | none |
| 125 | I | 5VDC | 47°C | 1587 | – | – | bright | bright | bright | slight | total | total | clear | none | none | none |
| 127 | J | 5VDC | 87°C | 1582 | – | – | bright | grey | bright | total | total | slight | yellowing | none | 0–5 | none |
| 128 | J | 5VDC | 87°C | 2 | 2 | wire bond o/c | sample removed; unavailable for visual inspection | | | | | | | | | |
| 129 | J | 5VDC | 87°C | 1582 | – | – | grey | grey | black | total | total+ | total+ | yellowing | none | 0–5 | none |
| 130 | J | 5VDC | 87°C | 1582 | – | – | black | black | bright | total | slight | total | clear | extensive (WE) | >5 | yes |
| 131 | J | 5VDC | 87°C | 1582 | ~1000 | intermittent o/c | grey | grey | grey | slight | none | slight | yellowing | none | >5 | none |
| 132 | J | 5VDC | 87°C | 45 | 45 | delamination at adaptor | sample removed; unavailable for visual inspection | | | | | | | | | |
| 134 | J | 5VDC | 87°C | 1582 | – | – | black | black | bright | total | total+ | total | yellowing | none | >5 | none |
| 135 | J | 5VDC | 87°C | 1582 | – | – | black | grey | grey | total | total+ | total | clear | extensive (CE, WE) | 0–5 | none |
| 136 | J | 5VDC | 87°C | 1582 | 1582 | bung leak | grey | black | black | total | total+ | total | clear | total (CE, SH, WE) | none | yes |
| 137 | J | 5VDC | 87°C | 500* | 0 | intermittent o/c | sample removed at Day 500; unavailable for visual inspection | | | | | | | | | |
| 138 | J | 5VDC | 87°C | 500* | 0–7 | intermittent o/c | sample removed at Day 500; unavailable for visual inspection | | | | | | | | | |
| 139 | J | 5VDC | 87°C | 1582 | – | – | grey | black | bright | none | total+ | total+ | cloudy | extensive (CE) | 0–5 | none |

**Table S2:** Visual changes observed in the CMOS-IDC samples.

| SAMPLE INFORMATION | | | | | | | ELECTRICAL DATA | | |
|---|---|---|---|---|---|---|---|---|---|
| **ID** | **Batch** | **Voltage** | **Temp.** | **Retirement Day** | **Fault/Failure Day** | **Failure/Fault** | **EIS Stability** | **Nature of Instability** | **General EIS Observations** |
| 86 | G | ±5V | 67°C | 1286 | – | – | stable | – | – |
| 87 | G | ±5V | 67°C | 1286 | – | – | stable | – | – |
| 88 | G | ±5V | 67°C | 1286 | – | – | stable | – | – |
| 89 | G | ±5V | 67°C | 1286 | – | – | stable | – | – |
| 90 | G | ±5V | 67°C | 1286 | – | – | stable | – | – |
| 91 | G | ±5V | 67°C | 0 | 0 | wire bond o/c | infant failure | – | – |
| 93 | G | ±5V | 67°C | 1286 | – | – | stable | – | \|Z\| increase at f > 10kHz |
| 94 | G | ±5V | 67°C | 1286 | – | – | stable | – | worm-hole void (badly-cured silicone) |
| 95 | G | ±5V | 67°C | 1286 | – | – | stable | – | badly-cured silicone |
| 99 | H | 5VDC | 67°C | 1419 | – | – | stable | – | – |
| 100 | H | 5VDC | 67°C | 1419 | – | – | stable | – | – |
| 101 | H | 5VDC | 67°C | 1419 | – | – | stable | – | – |
| 102 | H | 5VDC | 67°C | 1419 | – | – | stable | – | – |
| 103 | H | 5VDC | 67°C | 71 | 71 | wire bond o/c | unstable | open circuit | instability beginning at Day 34; then open circuit failure Day 71 |
| 104 | H | 5VDC | 67°C | 1419 | – | – | stable | – | $\theta$ fluctuations at f < 1Hz |
| 106 | H | 5VDC | 67°C | 71 | 71 | bung leak | unstable | SH-WE leak | removed after EIS artefact (SH-WE leak) |
| 107 | H | 5VDC | 67°C | 1419 | – | – | unstable | other | Intermittent increases in \|Z\| |
| 108 | H | 5VDC | 67°C | 441 | 441 | bung leak | unstable | other | highly unstable sample; failed at Day 441 from a short between the WE pin in the test tube bung and its surrounding SH tube |
| 109 | H | 5VDC | 67°C | 1419 | – | – | unstable | other | intermittent increases in \|Z\| |
| 110 | H | 5VDC | 67°C | 447 | 447 | wire bond o/c | unstable | open circuit | wire bond open circuit |
| 113 | I | 5VDC | 47°C | 1587 | – | – | stable | – | Temporary increase in \|Z\| around Day 1014 |
| 114 | I | 5VDC | 47°C | 1587 | – | – | stable | – | Temporary increase in \|Z\| around Day 1014 |
| 115 | I | 5VDC | 47°C | 1587 | – | – | stable | – | Temporary increase in \|Z\| around Day 1014 |
| 116 | I | 5VDC | 47°C | 1587 | – | – | stable | – | Temporary increase in \|Z\| around Day 1014 |
| 117 | I | 5VDC | 47°C | 1587 | – | – | stable | – | Temporary increase in \|Z\| around Day 1014 |
| 118 | I | 5VDC | 47°C | 1587 | – | – | stable | – | Temporary increase in \|Z\| around Day 1014 |
| 120 | I | 5VDC | 47°C | 1587 | – | – | stable | – | Highly stable |
| 121 | I | 5VDC | 47°C | 1587 | – | – | stable | – | Highly stable |
| 122 | I | 5VDC | 47°C | 1587 | – | – | stable | – | Highly stable |
| 123 | I | 5VDC | 47°C | 1587 | – | – | stable | – | Highly stable |
| 124 | I | 5VDC | 47°C | 1587 | – | – | stable | – | Highly stable |
| 125 | I | 5VDC | 47°C | 1587 | – | – | stable | – | Highly stable |
| 127 | J | 5VDC | 87°C | 1582 | – | – | stable | – | \|Z\| increase at f > 10kHz |
| 128 | J | 5VDC | 87°C | 2 | 2 | wire bond o/c | infant failure | – | – |
| 129 | J | 5VDC | 87°C | 1582 | – | – | unstable | other | \|Z\| & $\theta$ unstable at low & high frequencies |
| 130 | J | 5VDC | 87°C | 1582 | – | – | stable | – | Phase instability at high frequencies |
| 131 | J | 5VDC | 87°C | 1582 | ~1000 | intermittent o/c | unstable | open circuit | temporary period of open circuit (950 < Day < 1150); persistent high frequency open circuit |
| 132 | J | 5VDC | 87°C | 45 | 45 | delamination at adaptor | unstable | SH-CE leak | fault: SH-CE leak at thick film pads, intermittent starting at Day 28; failure: short SH-CE, Day 42 |
| 134 | J | 5VDC | 87°C | 1582 | – | – | stable | – | \|Z\| & $\theta$ unstable at low & high frequencies |
| 135 | J | 5VDC | 87°C | 1582 | – | – | stable | – | $\theta$ unstable at low frequencies |
| 136 | J | 5VDC | 87°C | 1582 | 1582 | bung leak | unstable | SH-WE leak | temporary SH-WE leak at Day 7 |
| 137 | J | 5VDC | 87°C | 500* | 0 | intermittent o/c | stable | – | initially open then healed |
| 138 | J | 5VDC | 87°C | 500* | 0-7 | intermittent o/c | unstable | open-circuit | open until Day 7, then healed |
| 139 | J | 5VDC | 87°C | 1582 | – | – | stable | – | \|Z\| increase at f > 10kHz & $\theta$ fluctuations |

**Table S3:** Electrical changes observed in the CMOS-IDC samples.

#### Supplement 2. Methods and Data Interpretation for EIS: Measurement Parameters, Spectral Artefacts, and Failure Analysis

This supplement serves three purposes:

(i) To provide the parameters set in the impedance analyser during the EIS measurements. This is essential information for anyone repeating the experiment.

(ii) To show how the impedance spectra are often distorted by unpredictable interference, but poor results can generally be rejected by examination of the records. Understanding this distortion improves one's confidence in the validity of the results.

(iii) To show how the recorded data were used in the investigation of failures, using ID106 and ID132 as examples.

**Instrumentation.** The ALTA apparatus was designed to perform long-term tests on the electrical insulation of candidate materials for implanted devices [1]. The test structures are planar interdigitated combs, coated or encapsulated, and maintained in fluid at elevated temperatures (in our work, up to 87°C). Electrical stress is applied most of the time, and deterioration is monitored by connecting the samples occasionally to an impedance analyser (interrupting the electrical stress). Each test sample is placed in its own individual test tube, and sets of tubes are heated by partial immersion in a water bath. In our apparatus, each hot water bath holds 96 tubes (hence, 96 samples).

This supplementary material assumes that the reader is familiar with the apparatus, its properties, and its performance, as described in [1].

The impedance analyser is a Modulab XM, developed by Solartron Analytical and sold by Ametek. It comprises three modules: a potentiostat, a femtoammeter, and a frequency response analyser (FRA). The potentiostat was designed for three-electrode measurements, using a working electrode (WE), a counter electrode (CE), and a reference electrode (RE). In the ALTA setup, the reference electrode connection is joined to the counter electrode connection, reducing it to a two-wire arrangement. The defined voltage is applied between the wires (RE-CE to WE), and the current is measured by a zero-resistance ammeter ("femtoammeter") at the WE. Multiplexing of the samples is described in [1].

**Frequency Response Analyser (FRA).** Used for Electrical Impedance Spectroscopy, the FRA measures the impedance at pre-defined frequencies (called $\omega_0$ in this paper), one by one. The FRA synthesizes the sinusoidal drive voltage ($V_{\mathrm{SET}}$ in Figures S1 and S2a), which is applied at CE. The current is amplified by the femtoammeter, which has adjustable gain, giving 12 ranges from 300 mA to 3 pA. Effectively, the FRA also generates a corresponding cosine waveform, and these two—sine and cosine—are multiplied by the voltage output of the femtoammeter and integrated over an integer number of cycles ($2\pi N$). The integrator outputs represent the real and imaginary parts of the complex current, from which the

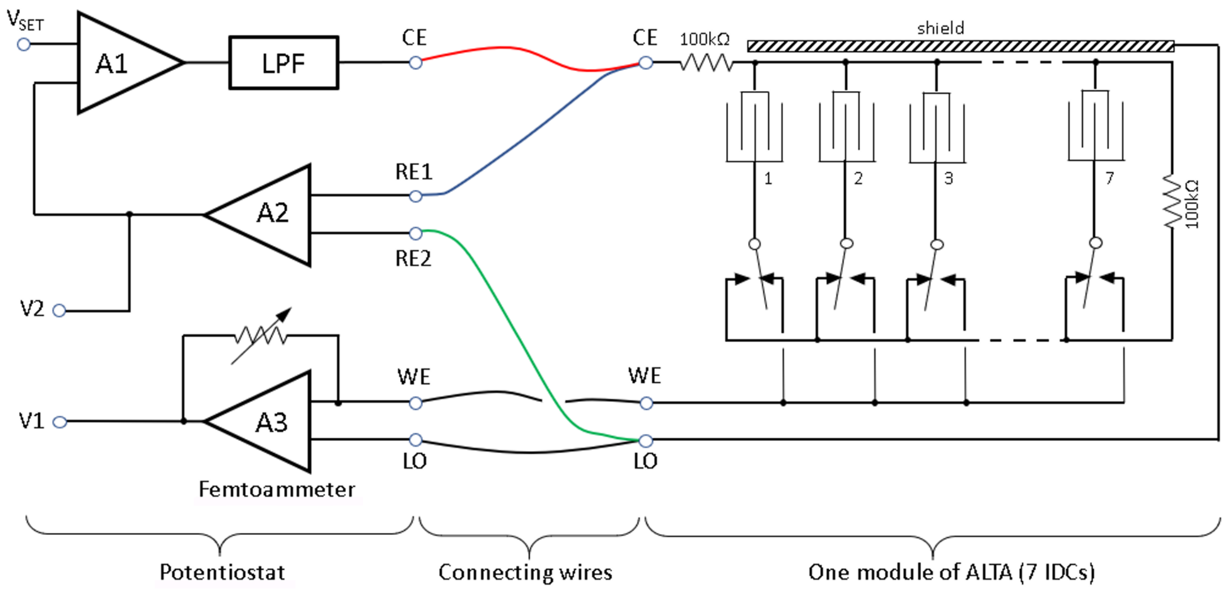


**Figure S1.** Connections between the potentiostat and the module of the ALTA being measured. See Donaldson (2018) [1], Figures 4 and 7; also Figure 6-2 in the *Modulab XM ECS User Guide*. RE1 is connected to CE, and RE2 to LO for two-wire impedance measurements. A3 is the femtoammeter. V1 and V2 are monitoring terminals on the Frequency Response Analyser (FRA) that allow the driving voltage and current to be observed with an oscilloscope (*Modulab Installation Guide*, page 1-15).

impedance is calculated (Figure S2a). This method was described by Gabrielli (1974) [2]; see McKubre and Macdonald (2005) [4].

In the Modulab XM FRA, multiplication and integration are performed by a digital signal processor at a rate as high as 300 kS/s.

The FRA acts as a frequency-domain filter, as shown in Figure S2b. For frequencies close to $\omega_0$, it acts like a narrow-band filter with a passband inversely proportional to $N$. For high frequencies, it acts as a low-pass filter with a cut-off rate of 90 dB/decade (for any $N$). This characteristic means that the FRA is good at removing high-frequency interference, particularly the mains frequencies (50 Hz and its harmonics), when measuring at frequencies below 1 Hz—where the effects of sample deterioration were expected to be seen [1]. It is still susceptible to spike interference, as can be seen in the spectra presented below.

**Data Files.** The FRA multiplies the measured current with sine and cosine waves and integrates the products at 300 kS/s to calculate the magnitude and phase of the impedance for each applied frequency. The data saved in a CSV file is relatively sparse to limit file sizes.

Each row of data includes the following:

- time to the second,
- voltage sample,

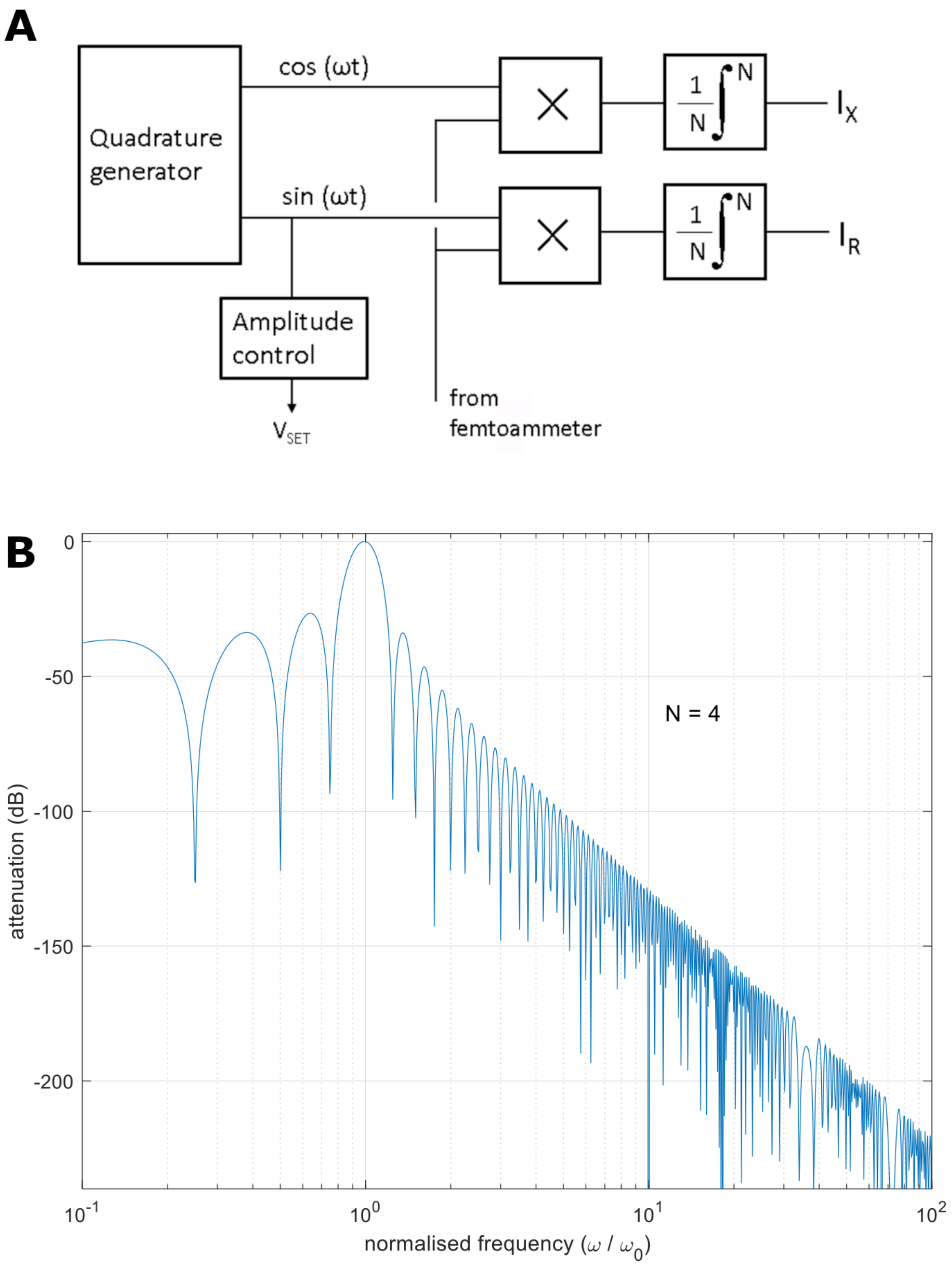


**Figure S2.** (a) Schematic diagram of FRA. (b) Frequency response of an FRA, integrating over 4 cycles at frequency $\omega_0$.

- current sample,
- frequency,
- impedance magnitude,
- impedance phase,
- voltage range,
- current range.

The first 24 rows give the impedances for the frequencies from 100,000 Hz to 2.5 Hz. From 1.6 Hz downward, there is a row every two seconds, but only the row at a time after a full integration (four cycles) contains impedance values. From this data, Bode plots can be drawn, and voltage and current waveforms can also be plotted. With samples only every two seconds, the Nyquist frequency is 0.25 Hz; above this, the plots are subject to aliasing.

**Table S4.** *Modulab "Experiment Setup" parameters. In our work, the instrument is set up for low-current electrochemistry* ($< 3$ mA*), which requires the potentiostat to have a sample rate of 1 MS/s and the FRA to have a computation rate of* 300 k*S/s.*

| **Parameter** | **Value** | **Comment** |
|---|---|---|
| Bandwidth | 100 kHz | Low-pass filter shown in Figure S1 |
| Voltage amplitude | 50 mV (r.m.s.) | |
| Current range max | 30 µA (range 5) | |
| Current range min | 300 pA (range 10) | |
| Auto-range speed | medium | |
| Frequency sweep | logarithmic | |
| Frequency sweep start | 100 kHz | |
| Frequency sweep stop | 10 mHz | |
| Frequencies per decade | 4 | 1.0 – 1.6 – 2.5 – 4.0 – 6.0 – 10 (approx.) |
| Integration time per frequency | the longer of: 0.1 s or 4 cycles | |
| DC measurement | periodic 2s/sample | Determines the sample rate in the data when the duration of four cycles is $> 2$ s |

**Examples.** Figure S3 shows four graphs presenting results from sample ID101 at Baseline (26/12/2017). The drive voltage is shown in (a). The coloured lines indicate the times when the frequency changed in the final decade, showing that integration occurred over four cycles at each frequency. The effect of aliasing can be seen in the apparent drive voltage amplitude, which appears smaller at 0.2 Hz and higher frequencies. The corresponding current is shown in (b). At the lowest frequency, the current is about $1 \times 10^{-12}\,A_{\text{p-p}}$, which means that the magnitude of the impedance is approximately $10^{11}\,\Omega$.

The Bode plots are shown in (c) and (d). This sample behaves like a capacitor at frequencies below approximately 1 kHz, with a capacitance of 156 pF. Its impedance magnitude reaches $10^{11}\,\Omega$ at 10 mHz, as expected. The Bode plots are only slightly affected by the phase deviating from quadrature (by 1.3°) at 10 mHz, due to interference visible in the current (Fig. S3b).

It is notable that at 1 Hz, the current range changed to 10 (300 pA), the lowest allowed value, yet no obvious quantisation appears in Figure S3b.

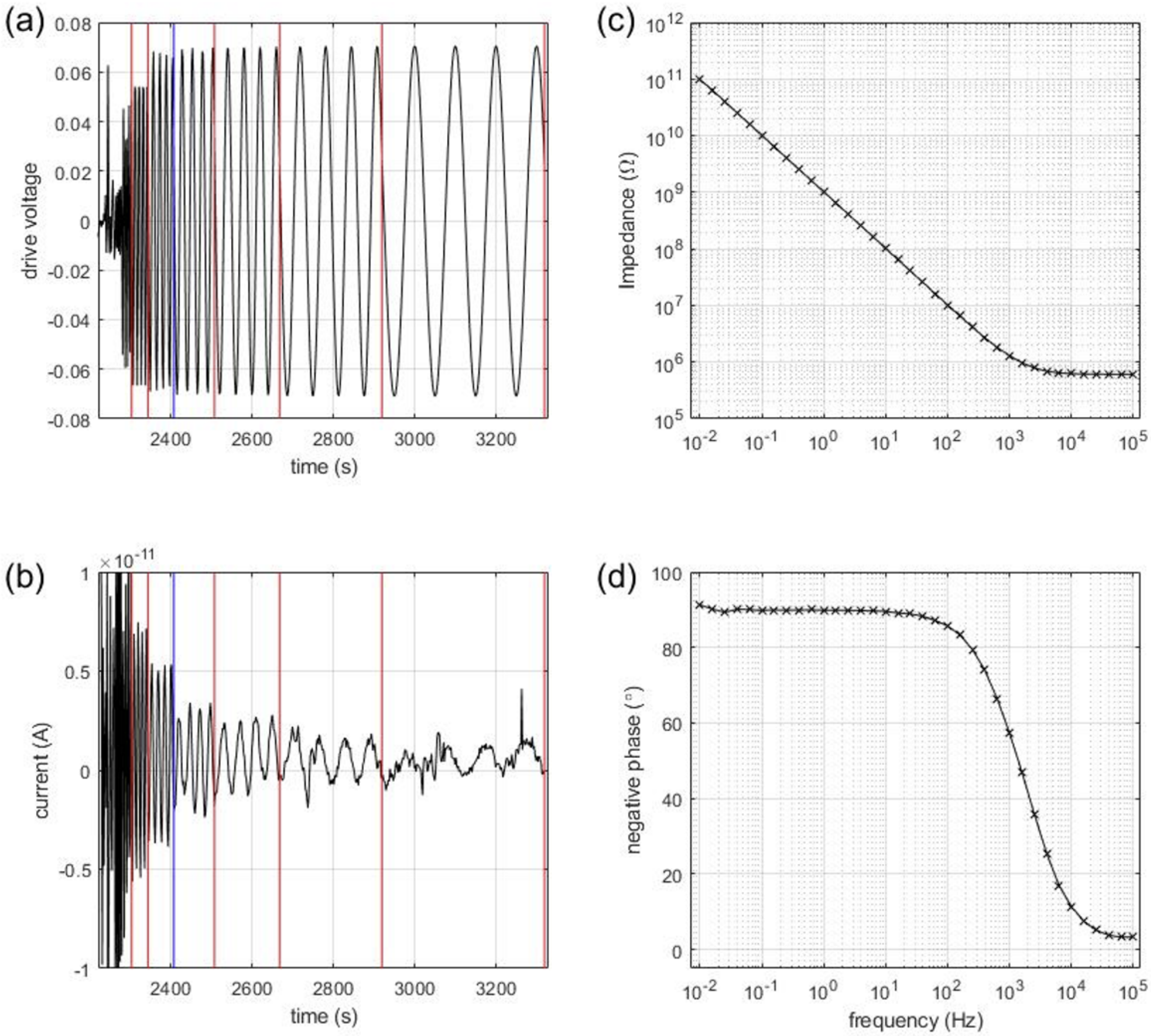


**Figure S3.** ID101 Baseline. The waveform is distorted by aliasing above 0.2 Hz, to the left of the blue lines in (a) and (b).

**Spectral Distortion at Low Frequencies.** The distortion of the Bode plots at the lowest frequencies, as seen in Figure S3, is common in our data. It typically appears first in the phase, which is more affected than the magnitude. There are three reasons why this distortion appears first at low frequencies:

- Given the capacitive nature of the samples—at least before any deterioration—the current decreases during the sweep, being proportional to frequency. Therefore, the amplitude of interference becomes relatively larger.
- The duration of the measurement at each frequency increases inversely with frequency,

making spike-like interference more likely at low frequencies. For example, if a single spike occurs during the measurement of a sample, the chance of it affecting the 10 mHz measurement is approximately 36%.

- Integration is performed over many more cycles at high frequencies than at low frequencies (see Integration Time in Table S4). At 100 kHz, there are 10,000 cycles in 0.1 s; at 100 Hz, there are 10; but below 40 Hz, there are only 4 or sometimes 5 cycles (adjusted by the experimenter).

The effect of random spike interference is consistent, as shown in Figure S4, which presents data from the same sample (ID101) after 1400 days at 67°C with a 5 V bias. There is a slightly larger spike in the current and a more pronounced distortion in the phase.

In addition to spike interference, Figure S4b shows periodic interference in the current during the last 600 s, apparently at about 45 mHz. However, due to possible aliasing, the true frequency of this interference cannot be determined.

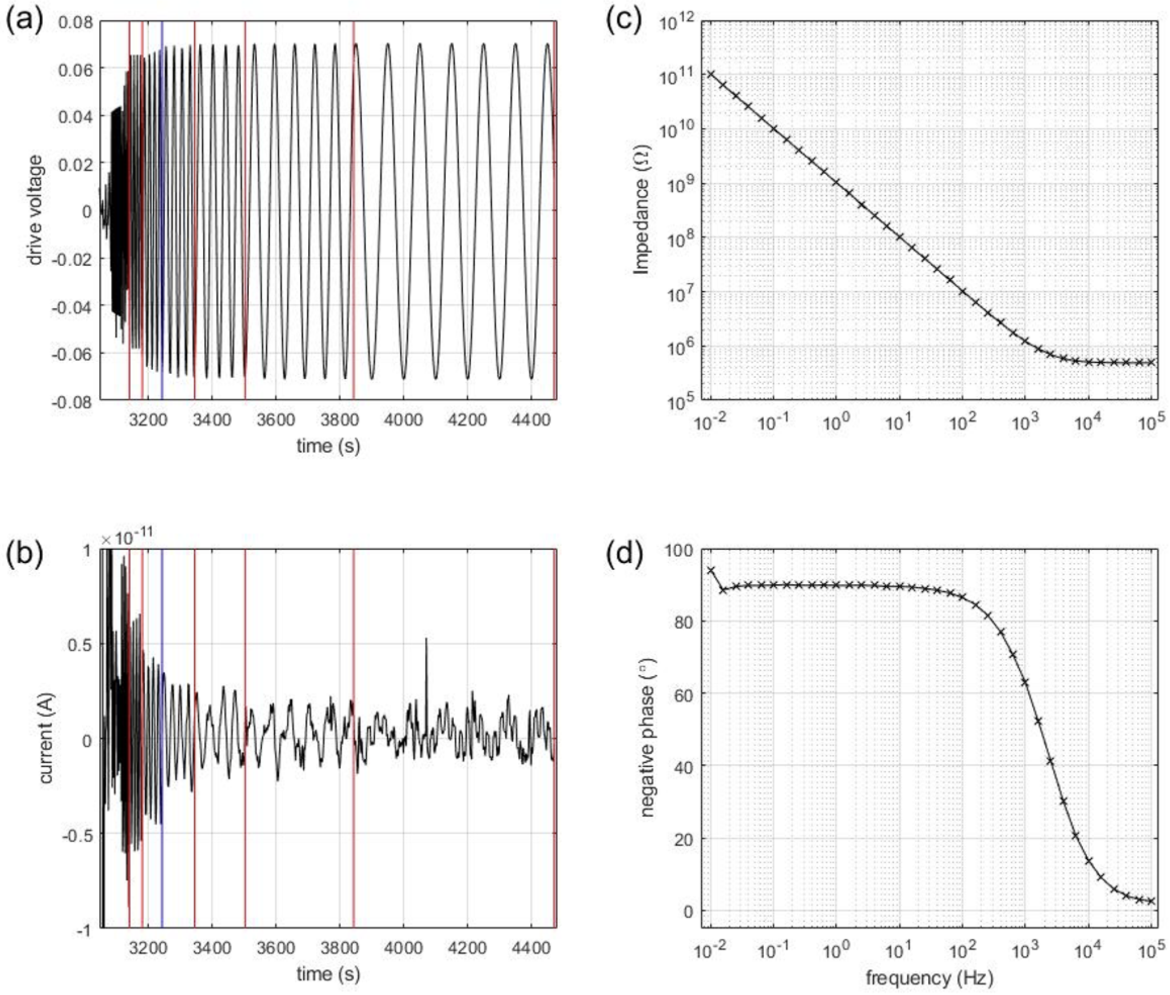


**Figure S4.** ID101 after 1400 days.

For samples that have failed open-circuit and for controls (tubes without sample), the impedance is even higher and therefore the results are more likely to display interference effects; for example, see Figure S5. In the current trace are many interfering spikes and the sinusoidal current is buried in the noise. Nevertheless, the impedance magnitude is only slightly distorted though the phase is badly distorted.

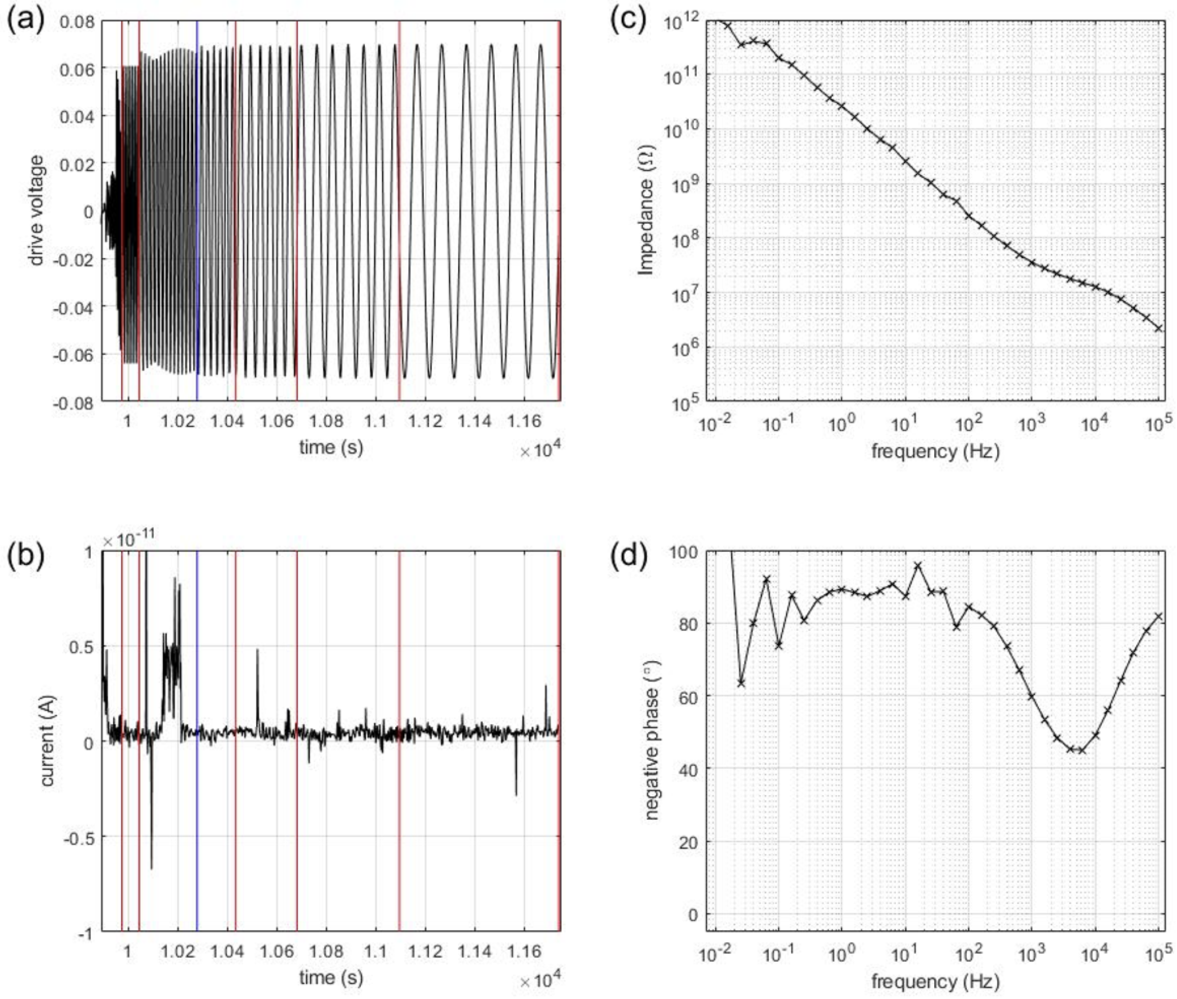


**Figure S5.** J14 on 10/5/2019 (Control in J Batch). Integration time has been increased to 6 cycles. At 10 Hz, $|Z| = 2.6\,\mathrm{G}\Omega$ which corresponds to a capacitance of 6 pF.

**Fault Revealed by Increased Impedance.** The magnitude of the impedance of sample ID132 had doubled by Day 42 (Figure 7, lower left panel). It was removed from the ALTA apparatus and, upon microscopic examination, was found to be delaminated at the adaptor, with the CE pad being the most affected. The impedance between CE and SH, measured with no other connections to the tube, exhibited constant-phase behaviour in the range 10–100 mHz (phase $\sim$55°), confirming that delamination had allowed leakage through water in the void.

In Figure S6 below, we present a photograph of this delamination in ID132. We report this to be the only sample in which silicone delamination was observed over the IC bond pads (at the SH and WE pads). The delaminations do not connect and thus are not detected electrically. The more obvious defect (and the one observed in EIS) was the CE-SH short on the ceramic adaptor.

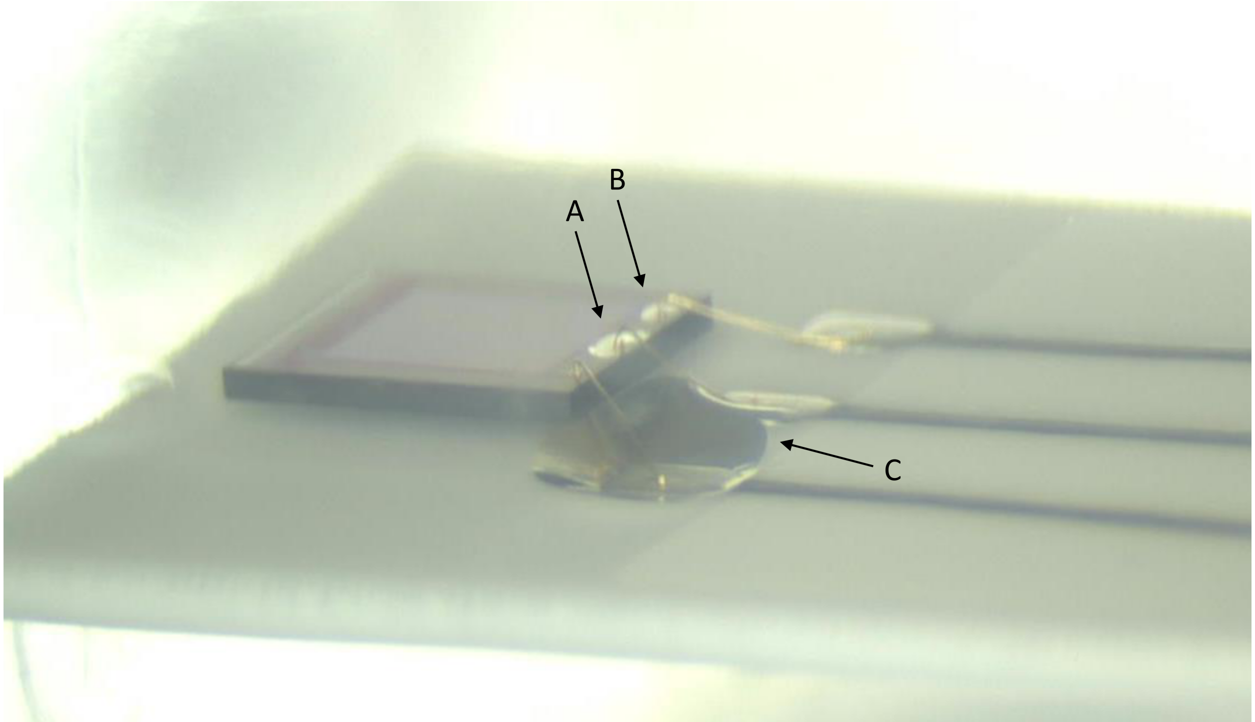


**Figure S6.** ID132, from Batch J (5V DC, 87°C). Delamination observed at Day 28 [3] and retired at Day 45. (A) Delamination of SH IC bond pad; (B) delamination over WE IC bond pad; and (C) delamination at the adaptor, with CE-SH short.

Why does a CE–SH leak *increase* the apparent impedance?

During EIS measurements (Figure S1), the circuit is complex but can be simplified as shown in Figure 16(b) of Donaldson (2018). The reduced circuit diagram comprises four impedances, as reproduced in Figure S6. At low frequencies (below 100 Hz), $R_1$ and $C_{\text{shunt}}$ are negligible, and the current flowing through the femtoammeter is determined by the reactance of the CE–WE capacitance of the IDC, $C_{\Delta WC}$.

However, if a leak develops from CE to SH that is not a much larger impedance than $R_1$, the voltage output from the potentiostat is attenuated at the CE node, reducing the current measured by the femtoammeter. This reduction in current appears as an *increase* in the EIS impedance (defined as the ratio of potentiostat voltage to femtoammeter current).

**Fault Revealed by Implausible Indicated Impedance.** On Day 71, the impedance spectrum for sample ID106 changed abruptly. It was normal down to 1 Hz but then deviated

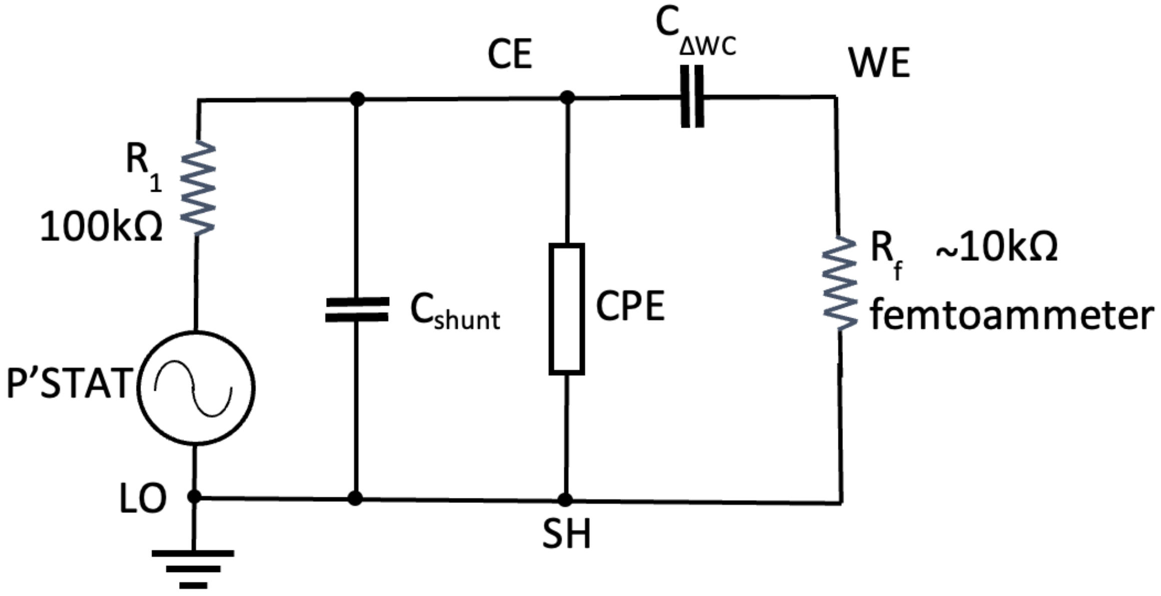


**Figure S7.** $C_{\text{shunt}} = C_{\Sigma}(\Delta C_S + \Delta W_S)$ in Figure 16b of Donaldson (2018) [1]. CPE is a Constant Phase Element representing the CE–SH water leakage.

dramatically from previous tests (Figure 7, lower right panel). Figures S7a and S7b show the corresponding time-domain record. The femtoammeter recorded a large, non-sinusoidal transient current. Given the magnitude of this artefact, it is remarkable that the FRA was able to measure the impedance accurately down to approximately 1 Hz; however, at lower frequencies—with fewer cycles available for integration—it appears to fail.

After removal from the ALTA apparatus, this sample was found to have a leakage impedance of only 70 MΩ below 1 Hz between SH and WE. This leakage was located at the bung, as it increased when the radial wire from the SH pin to the shield tube at the bung was cut (this wire is shown in Figure 5c in [1]).

**Discussion of Method.** When we planned this experiment, we expected that the samples would deteriorate much more quickly than they did, and so we aimed to test them frequently to monitor changes over time. As a result, we limited the time for each test, integrating over only four cycles (or occasionally six, as in Figure S5). However, the Modulab also allows integration to continue until the impedance at each frequency has converged to within either 10% or 1%, which should have reduced the interference effects described above. For samples that degrade as slowly as these, it would have been preferable to allow longer integration times and to measure impedance less frequently.

The CSV file for each EIS spectrum occupies only 45 kB. Although it does not have sufficient resolution to reveal the interfering frequencies (within the 200 kHz bandwidth), it does show when interference occurred during the low-frequency sweep. Consequently, Bode

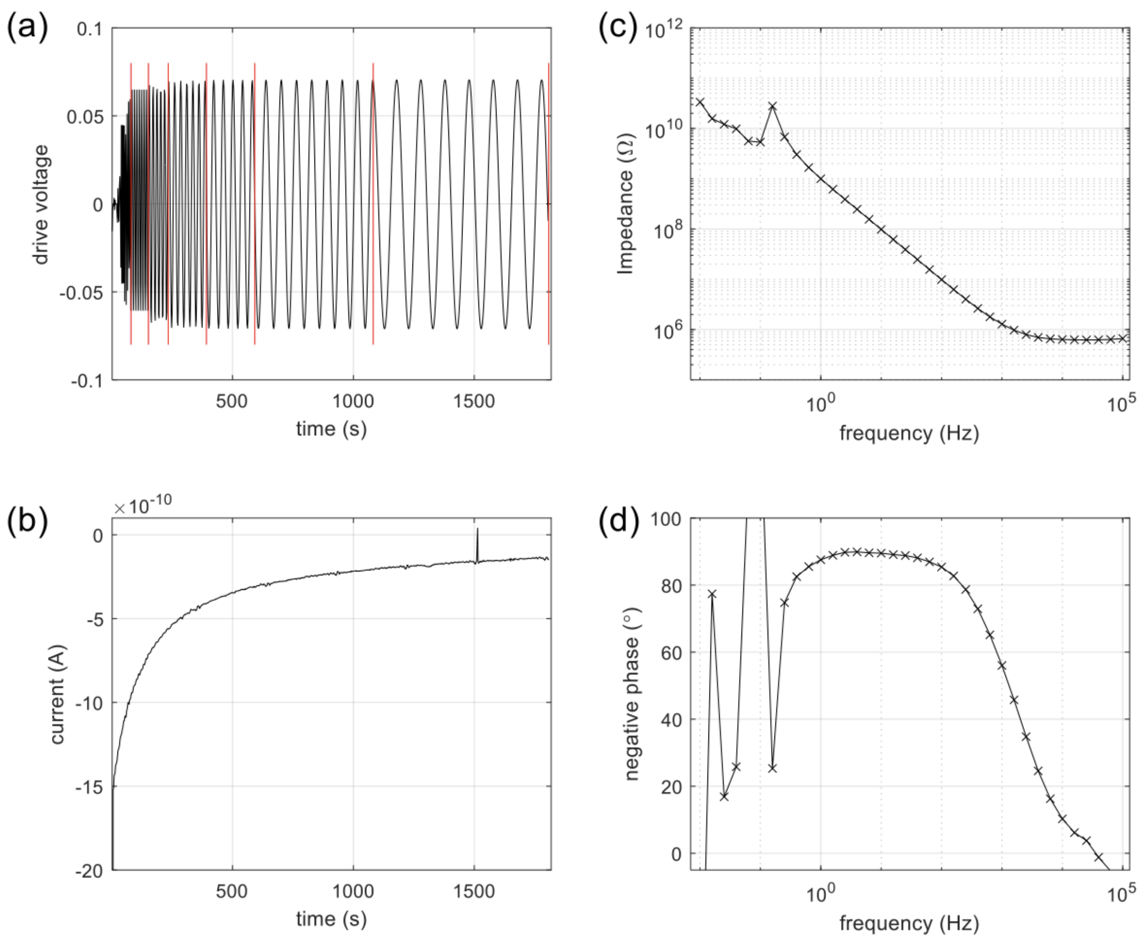


**Figure S8.** ID106 on Day 71.

plots should be interpreted with caution.

The CSV files in our experiments contained 643 lines. This could have been increased to 2000 lines (the Modulab limit) by increasing the data sampling rate, which would reduce aliasing. Another approach would be to reduce the frequency range—the high frequencies provided little useful information for our study—which would effectively reduce the bandwidth of the low-pass filter and thus the spike interference.

The examples shown in Figures S3 to S5 illustrate the limits of using the Modulab in combination with the ALTA apparatus in an electrically noisy London university setting. Good CMOS IDCs with capacitances around 150 pF produce visible currents of approximately $\pm 1$ pA. In contrast, for the control samples with capacitances around 6 pF, the sinusoidal current is often smaller than the background noise and interfering spikes. Occasionally, we observed a spike-free current trace and a corresponding impedance rising to about 3 TΩ.
